# Functional distinction between ionic and electric ephaptic effects on neuronal firing dynamics

**DOI:** 10.64898/2026.03.26.714388

**Authors:** Eirill Hauge, Marte J. Sætra, Gaute T. Einevoll, Geir Halnes

## Abstract

Neuronal activity alters extracellular ion concentrations and electric potentials. Ephaptic effects refer to the feedback influence that these extracellular changes can have on neuronal activity. While electric ephaptic effects occur on a fast timescale due to extracellular potential perturbations, ionic ephaptic effects are driven by slower, accumulative changes in ion concentrations. Among the previous computational studies of ephaptic effects, the vast majority have focused exclusively on electric effects, while ionic ephaptic effects have largely been neglected. In this work, we present an electrodiffusive computational framework consisting of two-compartment neurons that interact via a shared extracellular space. By accounting for both electric potentials and ion-concentration dynamics in a self-consistent manner, our framework enables us to explore the relative roles of electric and ionic ephaptic effects. Through numerical experiments, we demonstrate that ionic and electric ephaptic interactions play very different roles. While ionic ephaptic interactions increase population firing rates, electric ephaptic interactions primarily drive subtle shifts in spike timing. Furthermore, we show that these spike shifts cause the phase difference (the distance in spike times between a small collection of neurons) to converge to a stable, unique phase difference, which we coin the *ephaptic intrinsic phase preference*.

**Author summary:** Neurons predominantly communicate through synapses: specialized contact points where a brief electrical signal, known as a spike or action potential, in one neuron influences another. Neurons generate these spikes by exchanging ions with the surrounding extracellular space. This way, spiking neurons alter extracellular ion concentrations and electric potentials. Since neurons are sensitive to such changes in their environment, they can also influence one another indirectly through the shared extracellular medium. This form of non-synaptic interaction is known as *ephaptic coupling*. Most computational models of neuronal activity neglect ephaptic interactions, and those that include them typically consider only electric effects while ignoring ionic contributions. As a result, the relative roles of electric and ionic ephaptic effects remain poorly understood. Here, we introduce a computational framework that accounts for both mechanisms in a self-consistent way. Our results show a functional distinction: ionic ephaptic effects act slowly, regulating population firing rates, whereas electric ephaptic effects act on millisecond timescales and subtly shift spike timing. These shifts cause spike-time differences between neurons to converge to a stable value, a phenomenon we call *ephaptic intrinsic phase preference*.

## 1 Introduction

Ephaptic effects refer to indirect, non-synaptic influences that a neuron can exert on its neighbors (or itself) via activity-dependent modifications of its extracellular surroundings. In the literature, the term *ephaptic effects* most commonly refers to *electric ephaptic effects*, where extracellular fields generated by membrane currents in active neurons affect the membrane potentials of neighboring neurons. However, since neuronal transmembrane currents are mediated by ions, we may also have *ionic ephaptic effects* via activity-induced changes in extracellular ion concentrations [1, 2].

Modeling studies have suggested that electric ephaptic effects can synchronize spiking activity at the population level [3, 4]. At a smaller spatial scale, it has been demonstrated that ephaptic effects can be prominent in axon bundles [5–8], that ephaptic interactions can affect coincidence detection of synaptic inputs [9], and that ephaptic interactions between the soma of one neuron and a nearby axon or dendrite can be quite strong [10].

Ionic ephaptic effects have been less well studied, likely because variations in extracellular ion concentrations are believed to be quite small during normal neuron signaling. However, during pathological conditions such as epilepsy, stroke, or spreading depression, and also during non-pathological periods of intense neural activity, both intra- and extracellular concentrations have been shown to vary quite dramatically [11]. The effects that intra- and extracellular concentration changes can have on neuronal firing properties have been the topic of many modeling studies, but mainly using simplified single-neuron models and without making references to ephaptic coupling [12–17].

Since extracellular potentials are effectively instantaneous functions of transmembrane currents, electrical ephaptic effects take place on the same millisecond timescale as neural signaling. In contrast, ionic concentrations tend to vary on a slower timescale of seconds to minutes. The ionic ephaptic effects depend on accumulative concentration changes occurring during longer periods of neural signaling.

Electric ephaptic effects are conceptually simple. They are explained directly by the changes in the membrane potential *ϕ*_m_ = *ϕ*_i_ − *ϕ*_e_ (where the subscripts “m”, “i”, and “e” stand for membrane, intracellular, and extracellular, respectively) that occur when the extracellular potential *ϕ*_e_ changes. Membrane-current induced changes in *ϕ*_e_ can be predicted by standard volume-conductor theory [2].

In contrast, ionic ephaptic effects can be twofold. Firstly, extracellular concentration changes directly impact neuronal reversal potentials (see Eq. (17)), leading to changes in neuronal firing properties. Secondly, localized extracellular concentration shifts can lead to extracellular concentration gradients. These will, in turn, be accompanied by diffusion potentials in the extracellular space (ECS) [18–20], which will give a contribution to *ϕ*_e_ that is not accounted for by standard volume-conductor theory.

The relative contributions and roles of electric versus ionic ephaptic effects have not been thoroughly investigated, probably because their electrodiffusive nature makes it computationally expensive for modeling studies [2].

Here, we propose and use a modeling framework for exploring both electric and ionic ephaptic effects in a self-consistent manner. By self-consistent, we in this context mean that we keep track of all ionic concentrations in all intra- and extracellular compartments and ensure that there is always and everywhere a consistent relationship between ionic concentrations, the charge associated with them, and the local electric potential. The model is further introduced in Section 2.1 and described in detail in the methods section (Section 4).

Simulations of our model suggest that ionic ephaptic effects can be potent modulators of population-level mean firing rates. In contrast, electric ephaptic effects were found to have a negligible impact on average firing rates. However, we saw that electric ephaptic effects influenced the spike times of neighboring neurons relative to one another, consistent with previous studies suggesting a role for electrical ephaptic interactions in the synchronization of neural activity [21].

Interestingly, our investigations of the electric ephaptic effects led to the discovery of a, to our knowledge, novel phenomenon. When small groups of neurons were stimulated with constant and identical inputs (but with different stimulus onsets, such that the neurons did not fire simultaneously), electric ephaptic coupling caused the neurons to converge to a unique and stable phase difference. We refer to this phenomenon as *ephaptic intrinsic phase preference*, as we find the final, stable phase difference to be independent of initial conditions and hence appears to be an intrinsic system property.

## 2 Results

Presenting our results, we start in Section 2.1 by briefly introducing the self-consistent framework for modeling electric and ionic ephaptic coupling. This framework allows us to (a) strengthen or weaken all ephaptic effects simply by varying the size of the extracellular space, and (b) turn on or off ionic ephaptic effects by letting the neurodynamics be dependent or independent, respectively, of extracellular ion concentrations. Using this, we explore the relative importance of electric and ionic ephaptic effects on population firing rates in Section 2.2, and demonstrate and examine the novel phenomenon of *ephaptic intrinsic phase preference* in Sections 2.3 and 2.4.

### 2.1 A self-consistent model for electric and ionic ephaptic coupling

The self-consistent model for electric and ionic ephaptic effects is depicted in Fig. 1. It allows an arbitrary number of two-compartment neurons, consisting of one soma and one dendrite compartment, to interact ephaptically via a shared two-compartment ECS. Because extracellular electric fields arise from spatial separation of transmembrane currents, a two-compartment model is the minimal requirement for modeling extracellular fields and thus electric ephaptic effects. In the default model setting, the intra- and extracellular compartment volumes were specified such that the total intra- to extracellular volume ratio had a realistic value of 2:1.

**Fig 1.**
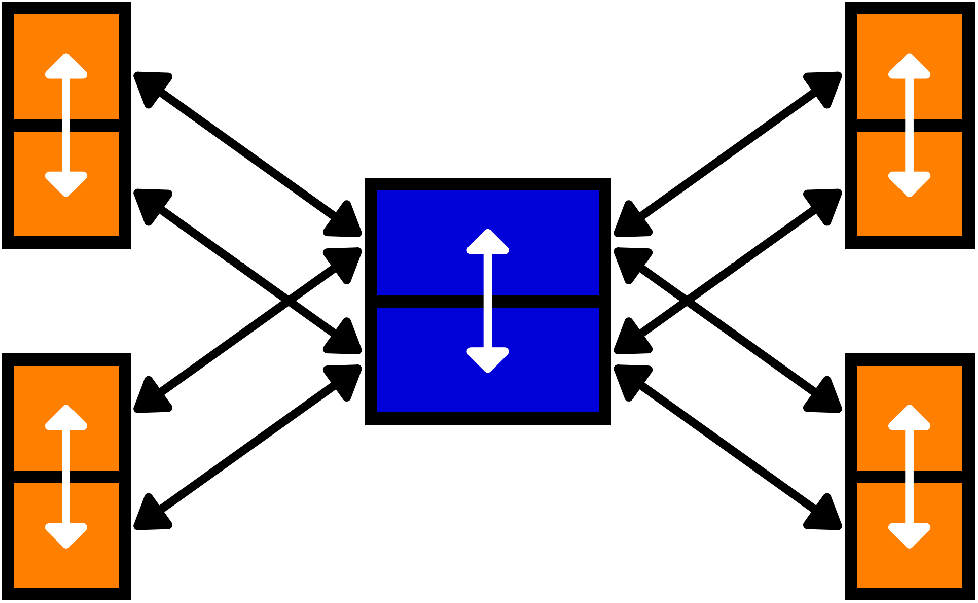
Model illustration. Four two-compartment neurons (orange) sharing a two-compartment ECS (blue). Each neuron has a soma-compartment (lower compartment) and a dendrite-compartment (upper compartment). Transmembrane currents (black arrows) in somas and dendrites enter or leave soma-level and dendrite-level ECS compartments, respectively. Transmembrane fluxes are modeled through Hodgkin-Huxley-type ion channels and additional pumps and co-transporters. Intra- and extracellular fluxes (white arrows) are modeled using the electrodiffusive KNP framework.

In the default setting, we used the two-compartment Pinsky-Rinzel model [22], as implemented in Sætra et al. (2020) [17], to model the neurons (see Methods Section 4.3). To test that our results can generalize to other models, we ran additional simulations using Hodgkin-Huxley mechanisms [23] for the membrane mechanisms (details are found in S1 Text). We will refer to the two different models as electrodiffusive Pinsky-Rinzel (edPR) and electrodiffusive Hodgkin-Huxley (edHH).

The dynamics of ions and voltages in all compartments were simulated using an electrodiffusive Kirchhoff-Nernst-Planck (KNP) framework [18, 19], which ensures a consistent relationship between ion concentrations (Na^+^, K^+^, Cl^−^, Ca^2+^, and an immobile generic anion X^−^ ), and electric potentials in all compartments at all times (see Methods Section 4.2).

The modeling framework allowed us to vary the magnitude of the ephaptic effects either by (i) varying the extracellular volume, or (ii) setting membrane currents to be dependent or independent of extracellular ion concentrations. By (i) increasing the extracellular volume, we could weaken both electric and ionic ephaptic effects (see Methods Section 4.5.1). By (ii) setting membrane currents (reversal potentials and pump parameters) to be independent of extracellular concentration changes, we could turn off the ionic ephaptic effects while keeping the electric effects (see Methods Section 4.5.2). We note that it was not possible to remove the electric ephaptic effects while keeping the ionic without compromising the biophysical consistency of the framework (see Section 3.1.3).

### 2.2 Influence of ephaptic coupling on population firing rates

To examine ephaptic effects, we studied changes in the population average properties of ten neurons as a function of the ECS size. All ten neurons received a distinct Gaussian-noise stimulus current with a mean strength ⟨*I*_stim_⟩ and 25% standard deviation (i.e., with standard deviation of 0.25 ⟨*I*_stim_⟩). The added stochasticity prevented the neurons from firing simultaneously. Using the same seed in each simulation, we ensured that each neuron received the same pseudo-random Gaussian current regardless of the ECS size. This way, changes in system behavior when varying the ECS size were not due to stochastic effects.

#### 2.2.1 Ephaptic coupling increases firing rates

To illustrate ephaptic effects in “raw data”, we compared spike trains between systems with regular-sized ECS and equivalent systems with the ECS size increased by a factor 10^6^ to weaken all ephaptic effects. We did this using three different stimulus strengths, with mean currents ⟨*I*_stim_⟩ of 16 pA, 28 pA, and 40 pA. Across all three stimulus strengths, the number of action potentials was visibly reduced when increasing the ECS size (right column Fig. 2) compared to the reference system with a regular-sized ECS (left column Fig. 2).

**Fig 2.**
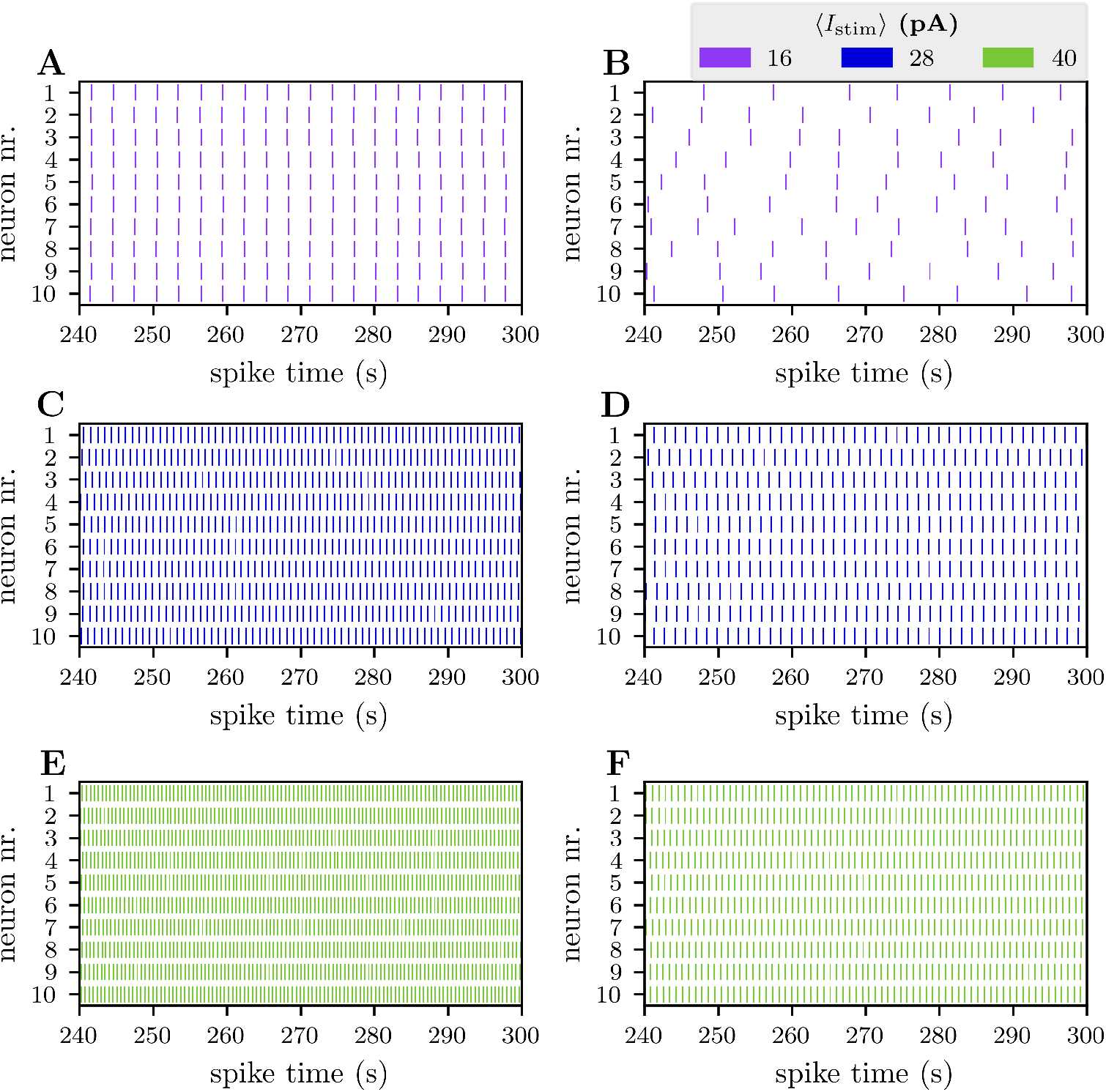
Spike times of 10 neurons during the fifth minute of simulation. All panels (**A**–**F**) show the spike times of 10 neurons, each receiving a noisy stimulus current with a mean value ⟨*I*_stim_⟩. Simulations presented in the left column (**A, C**, and **E**) were run with a regular-sized ECS, while the ECS was 10^6^ times larger in the simulations presented in the right column (**B, D**, and **F**). The ECS size was increased in the right column to weaken all ephaptic effects.

In general, the firing rate varied throughout the simulation, but settled on an approximately constant value after an initial (∼ 1 minute) transient following the stimulus onset (Fig. 3A). Specifically, with a mean stimulus ⟨*I*_stim_⟩ of 16 pA, 28 pA, and 40 pA, the systems settled at firing rates of respectively 0.33 Hz, 1.07 Hz, and 1.85 Hz. The ionic exchange associated with neuronal action-potential activity caused both intra- and extracellular ion concentrations to vary. Most notably, it led to an increase in the extracellular K^+^ concentration at the soma depth, 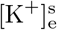. Also 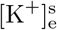 settled on an approximately constant value after an initial transient (Fig. 3C).

**Fig 3.**
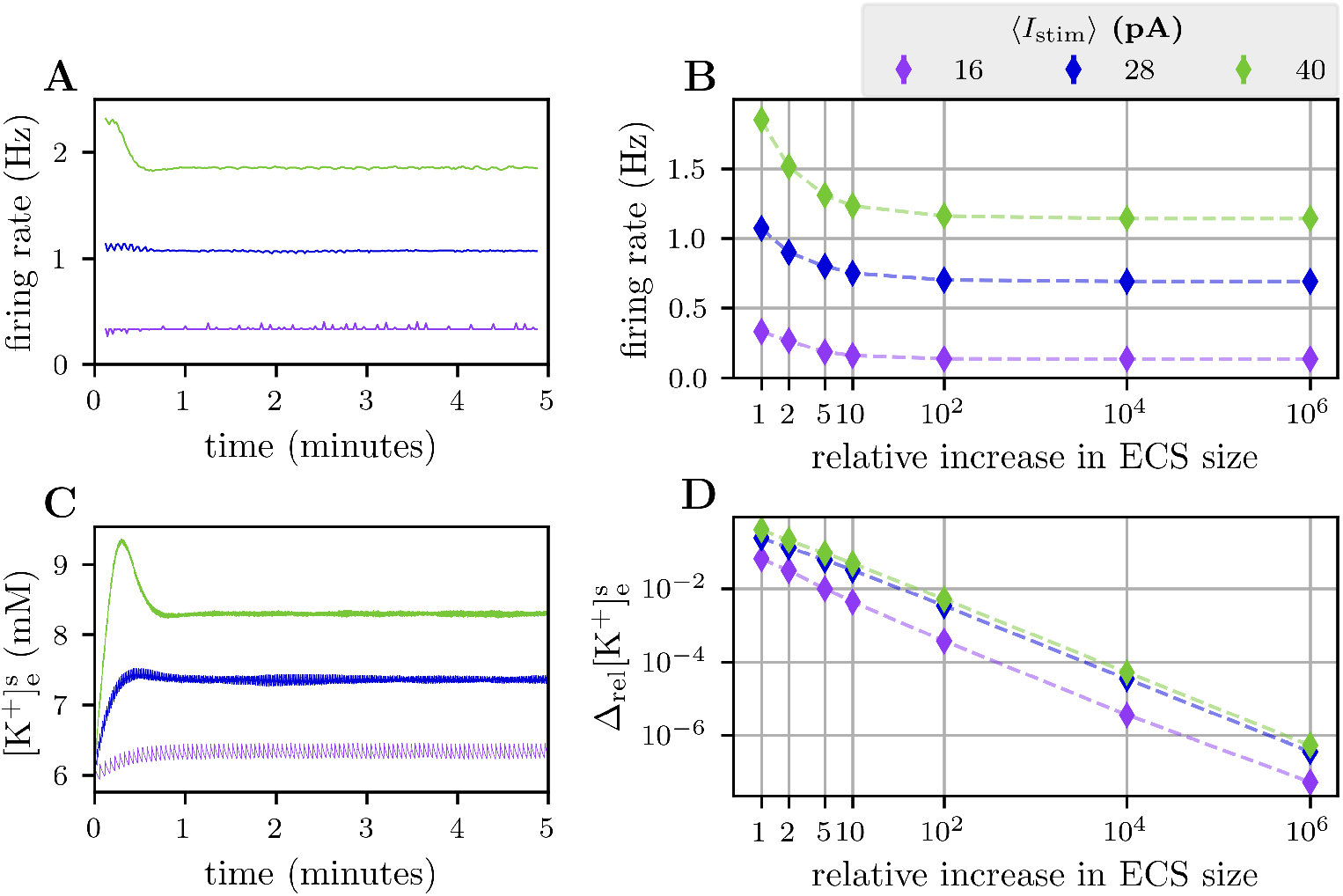
Population properties of ten neurons as a function of ECS size. Panels **A**–**B** depict the population-average firing rates as a function of time (**A**) and of ECS size (**B**). Firing rates in panel **A** were averaged using a sliding window of 15 s, and the systems had a default ECS size. Population averages in panel **B** were calculated over the fifth minute. Panels **C**–**D** illustrate changes in the extracellular potassium concentration at the depth of the somas 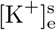 as a function of time (**C**) and of ECS size (**D**). All systems in panel **C** had a default ECS size. Panel **D** shows the relative deviation in 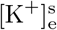, here defined as 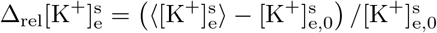, where 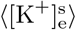 is the average of 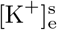 over the fifth minute and 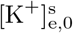 is the equilibrium concentration. In all simulations, all ten neurons received a Gaussian-noise stimulus current with a mean value ⟨*I*_stim_⟩.

Figure 3B and D summarize a number of simulations, and show how the *final* firing rate and 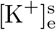 depended on the ECS size when we increased it from its regular size and up to a factor 10^6^ larger. As the figure shows, weakening the ephaptic effects by increasing the ECS size consistently decreased the final values for the firing rates (Fig. 3B) and 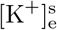 (Fig. 3D).

Increasing the mean stimulus strength ⟨*I*_stim_⟩ generally increased both the firing rate (Figs. 2 and 3A-B) and the extracellular K^+^ concentration (Fig. 3C-D). As previously stated, enlarging the ECS reduced the firing rates and limited the increase in 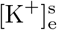 caused by neural activity. This trend of decrease in both firing rates and deviation of 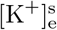, as a function of the ECS size, was observed for all stimulus strengths.

Over a large range in ECS size (from 1 to 100 times increase from the default ECS size), the size had a large impact on the firing rates and 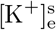. When the ECS size was increased beyond this, further ECS increases had only a minor impact on the firing rate, suggesting that ephaptic effects were becoming negligible for such large ECS compartments. Increasing the ECS from 10^4^ to 10^6^ times the default size did not give any changes in the firing rates (Fig. 3B). For both these ECS sizes, the firing rates remained stable at 0.13 Hz when ⟨*I*_stim_⟩ = 16 pA, 0.69 Hz when ⟨*I*_stim_⟩ = 28 pA, and 1.14 Hz when ⟨*I*_stim_⟩ = 34 pA.

We note that the relative change in extracellular 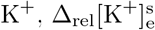, was close to inversely proportional to the ECS volume (declining linearly in the log-log plot) over the entire ECS-size range (Fig. 3D). Importantly, this does not imply the presence of ephaptic effects: increasing the ECS size simply reduces the concentration change associated with a number of ions entering this volume. For an ECS volume 10^6^ times the default value, 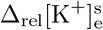 was less than 0.0001%, which we can assume to have a negligible impact on neuronal firing properties.

For use in the following studies, we conclude that increasing the ECS size by a factor 10^6^ (relative to the default value) is sufficient to eliminate all ephaptic effects.

#### 2.2.2 Increased firing rates are due to ionic ephaptic coupling

In the previous section (Section 2.2.1), we saw that ephaptic effects in general acted to increase the firing rates in our systems. Next, we wanted to investigate the relative roles of electric versus ionic ephaptic effects in causing this activity increase. To do this, we *turned off* ionic ephaptic coupling by setting the neuronal membrane dynamics (i.e., reversal potentials and ion pump rates) to be independent of extracellular ion concentrations. In this way, the neurons remained insensitive to changes in extracellular ion concentrations, but not to changes in the extracellular potential *ϕ*_e_. Hence, the firing rate was then affected by electric ephaptic coupling, but not by ionic ephaptic coupling.

As a reference, we used the same model setup as in Fig. 3, with ⟨*I*_stim_⟩ = 28 pA. In Section 2.2.1, we saw that the firing rate was reduced from 1.072 ±0.008 Hz to 0.692 ±0.008 Hz when we turned off all ephaptic effects by increasing the ECS volume by a factor 10^6^ (Fig. 3B). Note that the standard deviation measures the spread of firing rates among the ten neurons

To explore if this reduction in firing rate was explained by the loss of electric or ionic ephaptic effects, we reran the simulation with only ionic ephaptic effects turned off. A comparison of firing rates is illustrated in Fig. 4. Surprisingly, the firing rate for the system with only electric ephaptic effects (yellow: 0.690±0.008 Hz) was almost identical to the firing rate for the system with no ephaptic effects at all (orange: 0.692 ±0.008 Hz). This shows that electric ephaptic effects had almost no impact on the population firing rate. From this, we conclude that ephaptic effects on the population firing rates were almost exclusively due to ionic ephaptic coupling.

**Fig 4.**
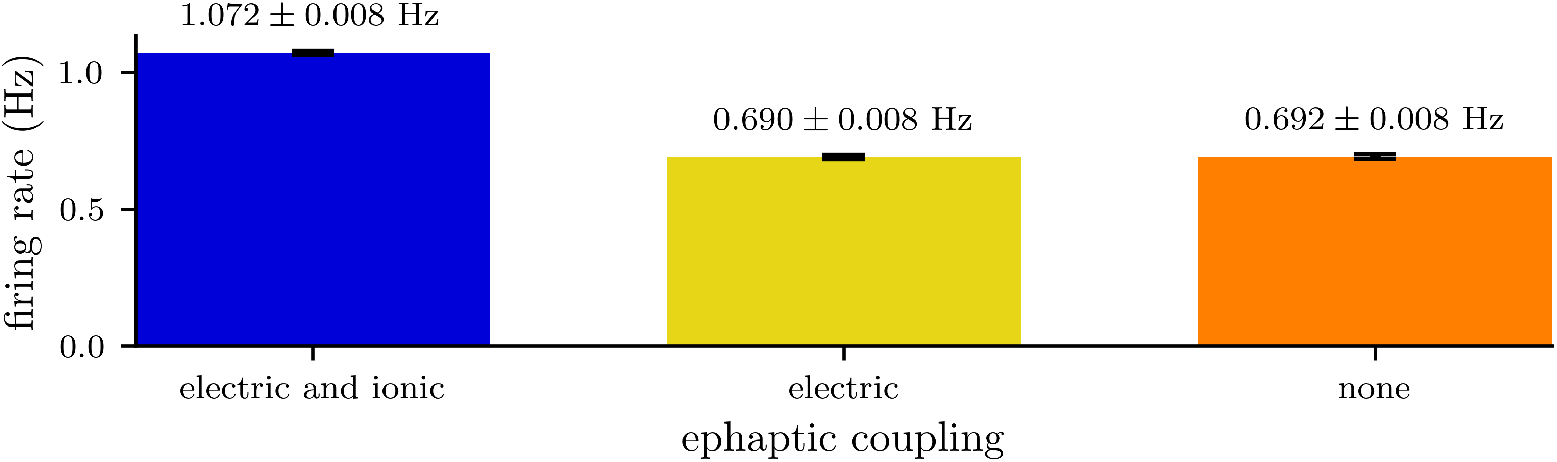
Firing rates of the same system with and without ephaptic effects. Comparison of firing rates between a system with both electric and ionic ephaptic effects (blue), electric ephaptic effects only (yellow), and no ephaptic effects (orange). Firing rates were calculated as the population average of the fifth minute in a system of ten neurons. Each neuron received a noisy stimulus with a mean strength of ⟨*I*_stim_⟩ = 28 pA. The standard deviation measures the spread of firing rates among the neurons.

This conclusion did not depend on our choice of neuron model. In simulations where we replaced the edPR model with the edHH model, the firing rates remained sensitive to ionic ephaptic effects, and almost unaffected by electric ephaptic effects (S1 Fig).

### 2.3 The importance of timing for ephaptic effects

All neurons in the system share the same extracellular space and therefore experience identical extracellular ion concentrations and potentials. If there are differences in how two neurons are (ephaptically) affected by the extracellular variables at a given time, it can only depend on differences between the individual neurons’ states at this time. For example, if a neuron is close to its firing threshold, it is likely more sensitive to perturbations.

In the current section, we are interested in studying how a single, regularly firing neuron is affected by an “ephaptic kick” associated with a neighboring neuron firing a single action potential. To do this, we consider a system of only two neurons.

We let the first neuron be a regularly spiking neuron, which received a constant stimulus 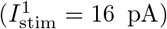 throughout all simulations. To obtain its regular firing state, a simulation was run where only this neuron was stimulated, while the other neuron received no input and remained silent. This simulation was run until the first neuron settled at a regular firing rate with a constant interspike interval (ISI). We used the second-to-last ISI as a reference ISI (area without shading in Fig. 5A), representing the ISI in the absence of ephaptic interactions with the other neuron.

**Fig 5.**
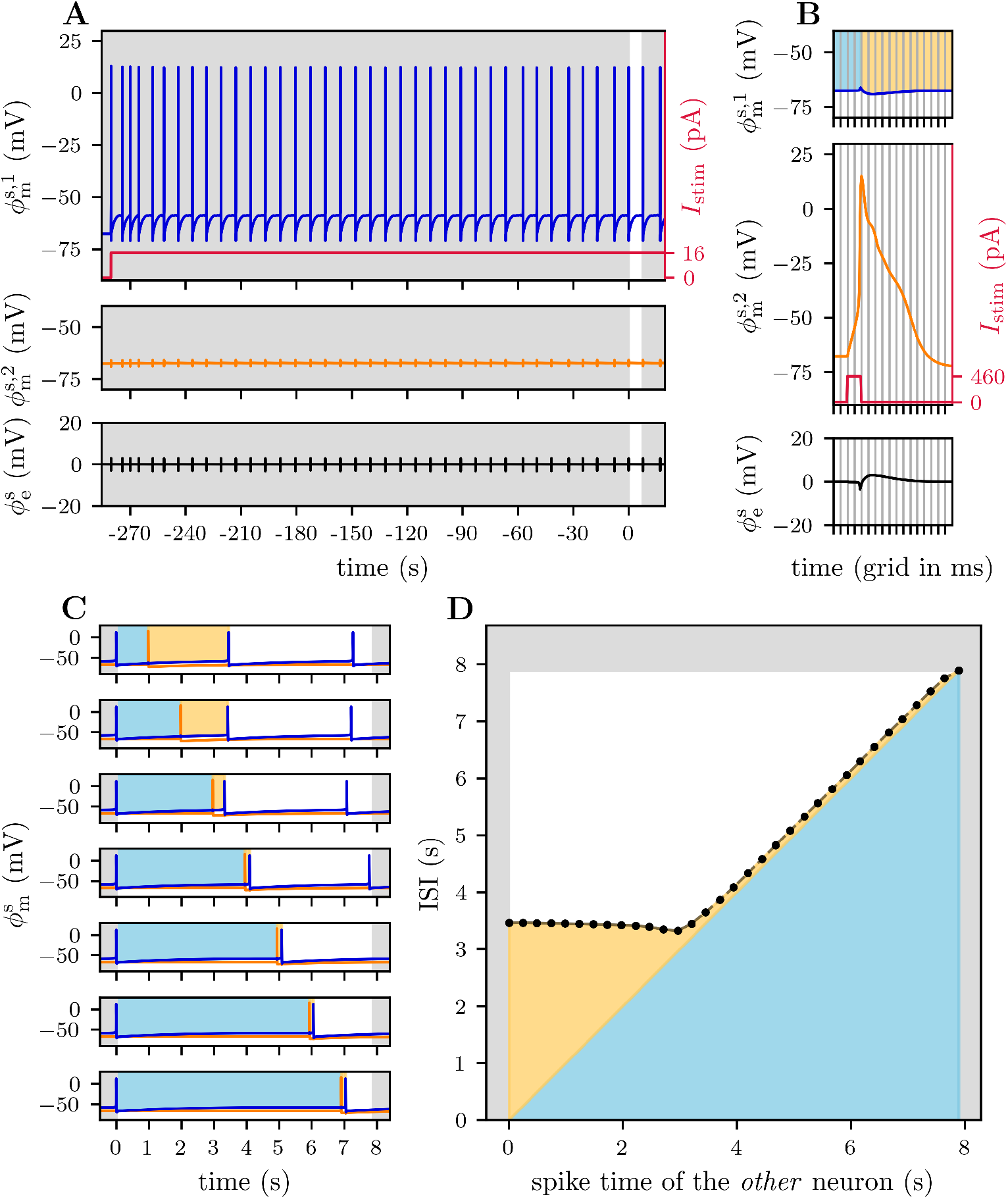
Impact of the timing of ephaptic coupling on the ISI. Panels **A** and **B** show the membrane potentials of the first neuron (upper plot) and second neuron (middle plot), as well as the extracellular electric potential (lower plot). Panel **A** shows the reference ISI (no gray shade) from a steady firing state where only the first neuron receives a stimulus current, while panel **B** illustrates how a stimulus pulse on the second neuron causes a single action potential, used in the simulations in Panels **C**–**D**. Note that the exact timing of the pulse varies within the reference ISI and is therefore not specified in the illustration. Panel **C** shows the membrane potentials of both the first neuron (blue line) and the second neuron (orange line). In each of the seven plots, the second neuron has a different spike time, affecting the ISI of the first neuron. Panel **D** shows the ISI of the first neuron as a function of the spike time of the second neuron. In panels **B**–**D**, the ISI of the first neuron is shaded light blue and yellow, corresponding to the intervals before and after the ephaptic kick (second neuron spikes), respectively.

Next, we introduced the ephaptic kick by triggering the second neuron with a brief and strong input pulse, making it fire a single action potential (Fig. 5B) at some time point within the reference ISI. We were interested in investigating how this kick affected the ISI, and to which degree this depended on the exact *timing* of this kick.

The stimulus current pulse to the second neuron was given by

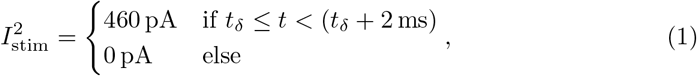

where the time of the pulse *t*_*δ*_ was chosen inside the reference ISI of the first neuron, indirectly controlling the spike time of the second neuron. To ease notation, we denote the (single) spike time of the second neuron as 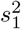, and label the spike times of the reference ISI as 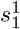 and 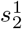, and shift the time such that 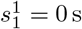.

By varying the spike time of the second neuron 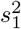, we compared the resulting length of the ISI of the first neuron with the reference ISI in Fig. 5A. Regardless of 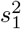, the single spike elicited by the second neuron shortened the ISI of the first neuron (Fig. 5C).

Further, we examined how *timing* of the ephaptic kick (by varying 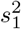) impacted *how much* the ISI was shortened (Fig. 5D). When 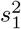 occurred in the later stages of the reference ISI 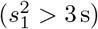, the regularly firing neuron was close to its firing threshold, and the ephaptic input made it fire shortly after 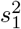. When 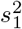 occurred in this later stage of the reference ISI, the time-lag between 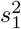 and the following spike 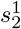 was nearly independent of 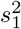. Hence, we observed a close-to-linear relationship between the ISI length and the timing of 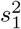 (Fig. 5D) in this regime.

When 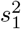 occurred in the earlier stages of the reference ISI 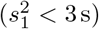, the trend was different. Although the time-lag between 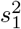 and the following spike 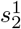 was longer in this case, it did lead to a significant shortening of the ISI (relative to the reference ISI). Interestingly, the ISI was a weakly declining function of 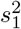 in this region, which reached a minimum around 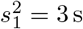 (Fig. 5D).

We conclude that (1) a neighboring neuron’s spike at a time 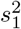 within the ISI of a regularly spiking neuron, shortens the ISI relative to what it would have been if ephaptic effects were absent, and (2) that there is an optimal time point 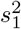 that results in the shortest possible ISI.

This conclusion was not strongly dependent on the applied stimulus to the regularly spiking neuron or the choice of neuron model. We arrived at the same qualitative conclusion using other stimuli (S2 Fig) and when we replaced the edPR model with the edHH model (S3 Fig). In one special case with the edHH model (4 pA stimulus), the optimal time 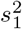 that resulted in the shortest possible ISI was found to be zero. That is, the shortest ISI was obtained when the ephaptic kick from the neighbor was perfectly synchronized with a spike of the regularly firing neuron.

### 2.4 Ephaptic effects on spike timing

In Section 2.3, we observed that the spike-time shift due to ephaptic effects depended on the *timing* of the ephaptic interaction. Importantly, the greatest impact on the neuron’s spike-time shift was achieved when it received an “ephaptic kick” at a specific delay from its previous spike time. Due to these observations, we wanted to investigate possible synchronizing effects and spike timing relationships between ephaptically coupled neurons.

To study possible changes in spike timing due to ephaptic effects, we considered *N* identical neurons that all received a constant (not noisy) stimulus current of the same strength, *I*_*c*_. Stimulus onsets to neurons *n* = 1, 2, …, *N* were applied sequentially, with a delay Δ_lag_ between the onsets of consecutive neurons:

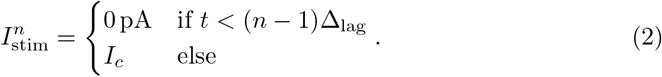

Here, Δ_lag_ will be referred to as the *initial delay*.

We define the *phase difference* in spike times as

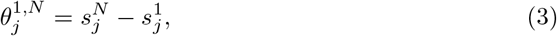

where 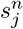 is the *j*’th spike time of the *n*’th neuron.

The only difference between the *N* neurons was the onset of their stimulus current, which in turn made them spike at different times. Note that the system was deterministic. Therefore, if the neurons had not been interacting ephaptically, we would expect them to spike in a sequence with constant delays Δ_lag_ between the spike times of consecutive neurons and a constant phase difference of 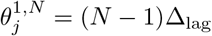.

#### 2.4.1 Ephaptic coupling creates a preferred phase shift between neurons

We started our study of spike-timing relationships using a simple system containing only two neurons, both receiving a 28 pA constant current injection. The first neuron received the current injection from *t* = 0, and the second neuron received the current injection from *t* = Δ_lag_. We explored three different cases, with initial delays of 10 ms (Fig. 6A), 50 ms (Fig. 6B) and 100 ms (Fig. 6C).

**Fig 6.**
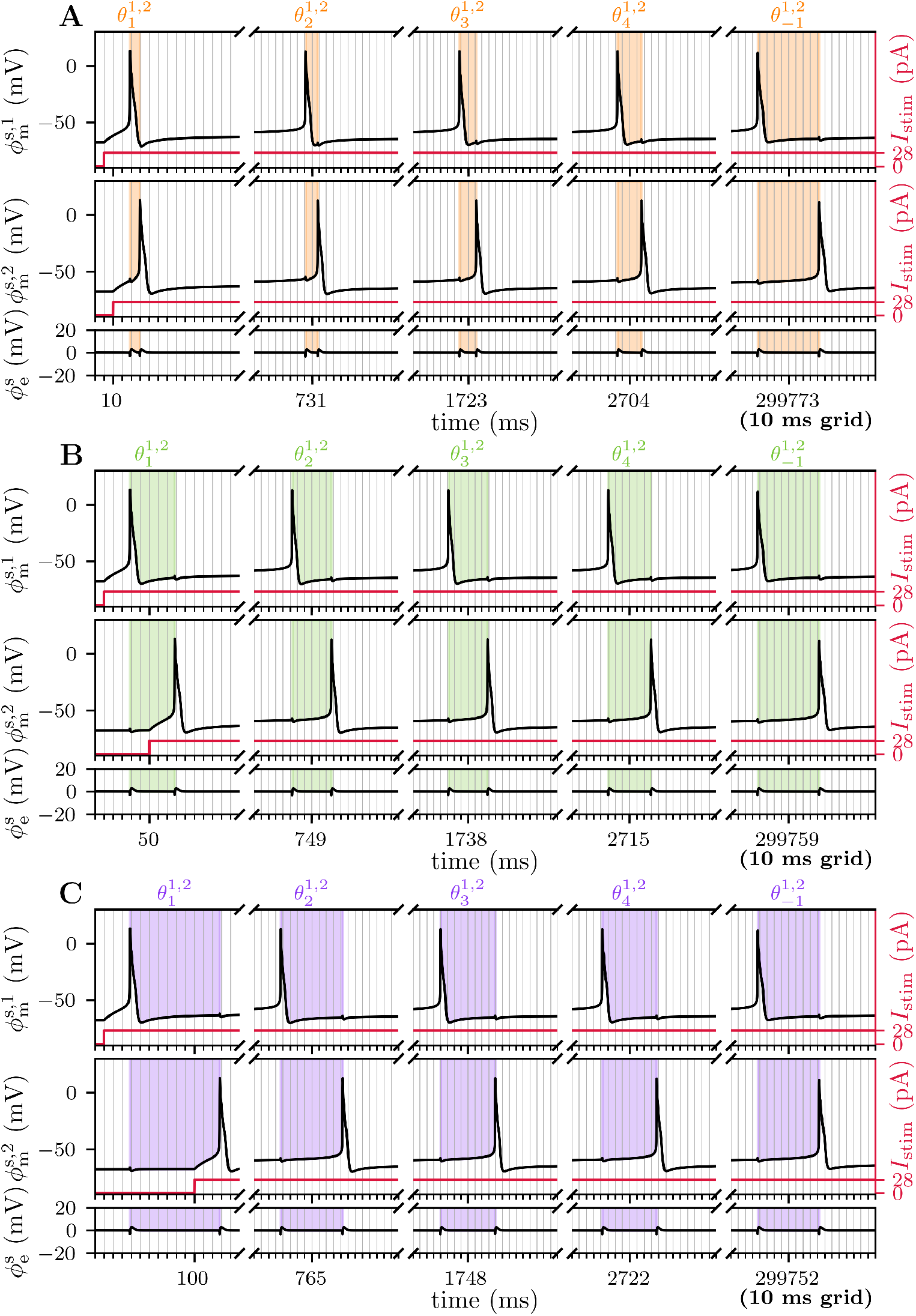
Tracking the spike phase 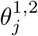 of two neurons over time. Each of the three panels shows the membrane potential of the first (upper plot) and second neuron (middle plot), as well as the extracellular potential (lower plot). Both neurons receive a constant stimulus current, where the onset of the second neuron’s stimulus current is delayed by Δ_lag_, set to 10 ms in **A**, 50 ms in **B**, and 100 ms in **C**. Note the use of a broken x-axis, each part spanning 160 ms. The distance between the *j*’th spike time of the first and second neuron, 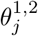, is shaded.

If the neurons had not been interacting ephaptically, they would end up firing with a constant phase difference

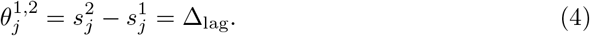

However, regardless of the specific initial delay value, we observed a dynamic phase difference (Fig. 6). This means that, similar to what we observed in Section 2.3, ephaptic interactions caused slight spike-time shifts. A dynamic phase difference implies that the magnitude of these spike-time shifts differs between the two neurons.

When using an initial delay Δ_lag_ of 10 ms, 50 ms, and 100 ms, the first phase difference 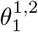 was 11.0 ms, 49.5 ms, and 99.5 ms, respectively. Hence, depending on timing, ephaptic effects could lead to both a prolonged (first case) or shortened (the last two cases) phase difference.

Note that the ephaptic interplay shifted the spike times of both neurons, so that the development of *θ*^1,2^ over time will depend on the *relative* change in the first versus second neuron. For example, if the spike times of the second neuron are accelerated more than the spike times in the first, *θ*^1,2^ will decrease. Conversely, if the spike times of the first neuron are accelerated more than the spike times of the second, *θ*^1,2^ will increase.

We ran the simulations for five minutes of simulated time. Initially, *θ*^1,2^ varied with time, but as the simulation progressed, the system converged to a stable firing state (1.03 Hz for both neurons). With a stable firing rate, the phase difference *θ*^1,2^ stabilized, remaining unchanged in the last minutes of simulated time (Fig. 7A). Surprisingly, the phase difference converged to the same value, regardless of initial conditions, that is, regardless of Δ_lag_ (Figs. 6 and 7A).

**Fig 7.**
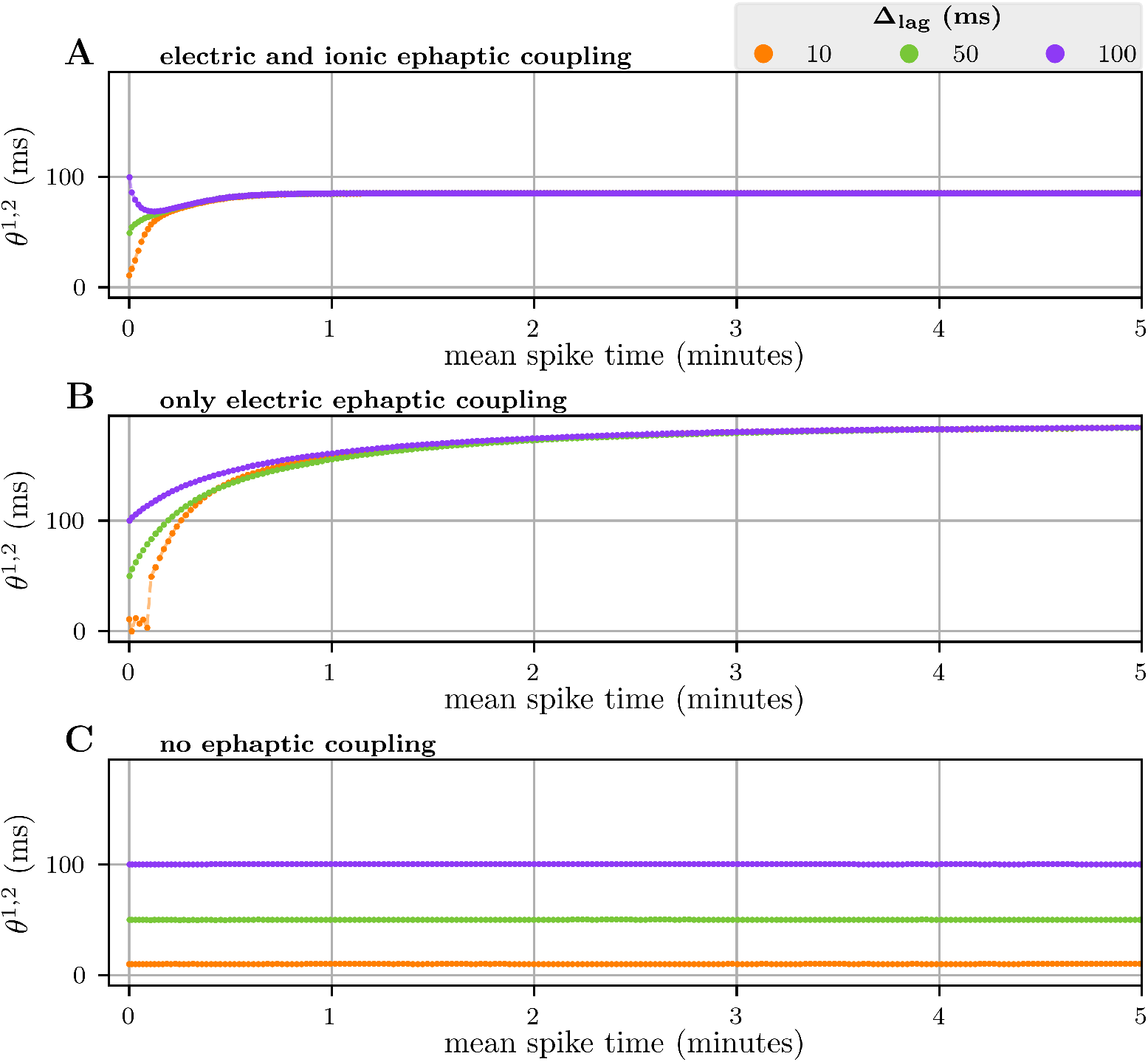
The spike phase with and without ephaptic coupling. Each panel shows the spike phase *θ*^1,2^ (the spike time difference between the first and second neuron) over time, for a system with both electric and ionic ephaptic coupling (**A**), electric ephaptic coupling only (**B**), and no ephaptic coupling (**C**). The system consists of two neurons, stimulated with a constant stimulus current of 28 pA and onset delay Δ_lag_ in the second neuron’s stimulus.

To quantify the converged spike-phase difference, we introduce the mean phase difference 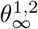, calculated as the average phase difference over the last minute of the simulation. For the case in question, the converged 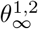 was calculated (over the fifth minute) to be 85.2 ms, 85.1 ms, and 85.1 ms, when using an initial delay Δ_lag_ of 10, 50, and 100 ms, respectively.

Both experimental [21, 24–27] and computational [3, 4, 10, 28, 29] studies have previously demonstrated how ephaptic (field-mediated) interactions can have synchronizing effects on neuronal firing. While related, the effect that we have demonstrated in this work is not synchronization in the classical sense, as they do not generally cause the neurons to fire more closely in time.

Our results are more closely related to the studies by Park et al. (2005) [30] and Stacy et al. (2015) [29], who both investigated interactions between a pair of neurons in a setup where ephaptic effects were accounted for. Both these studies found that electric ephaptic effects affected spike timing and influenced the phase-locking of neuron pairs. Our results extend these findings by demonstrating that ephaptic coupling can drive neurons to converge toward a preferred and stable phase difference that is independent of their initial phase relationship. As it appears to be an intrinsic property of the coupled system, we refer to it as *ephaptic intrinsic phase preference*.

Convergence of the phase to a constant value indicates that the ephaptic “kick” from neuron one advances the next spike of neuron two (relative to when it would occur without this kick) by exactly the same amount as the reciprocal ephaptic kick from neuron two advances the next spike of neuron one. That is, if the spike times of the two neurons are advanced by the same amount, the distance between them (the phase) will stay the same. One way in which this condition can be satisfied is if the interval from a spike in neuron one to a spike in neuron two equals the interval from the spike in neuron two to the subsequent spike in neuron one. This was not the case in our simulations. The other way in which this condition can be satisfied is if the neuron’s ISI as a function of kick-time exhibits at least one extremum, so that there are at least two kick moments that result in the same spike-advancement. We earlier saw that this was the case for our kick-time controlled simulations, where the ISI-curve had a clear minimum (Fig. 5D). The existence of an ephaptic intrinsic phase preference suggests that the same also holds for this more complex scenario of two continuously interacting neurons (see S2 Text for a more detailed discussion).

#### 2.4.2 Ephaptic intrinsic phase preference is due to electric ephaptic effects

The previous study (in Section 2.4.1) showed that the spike time distance between two neurons converged to an ephaptic intrinsic phase preference, 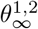. To examine what kind of ephaptic interactions are responsible for this phenomenon, we compared the time evolution of the spike-phase difference *θ*^1,2^ of three different cases with (i) both kinds of ephaptic effects present, (ii) only electric ephaptic effects present, and (iii) no ephaptic effects present. For all three cases we used the same two-neuron setup as in Fig. 6, and ran three simulations per case for different initial delays, Δ_lag_ ∈ {10 ms, 50 ms, 100 ms}. The results are shown in Fig. 7.

As expected, in the case (iii) with *all* ephaptic effects “turned off” (by enlarging the ECS by a factor 10^6^), *θ*^1,2^ remained constant and identical to the initial delay Δ_lag_ throughout the simulation (Fig. 7C). When ephaptic effects were present, *θ*^1,2^ varied with time, but converged to an ephaptic intrinsic phase preference 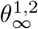 both in case (i) with both kinds of ephaptic effects present (Fig. 7A) and in case (ii) with only electric ephaptic effects present (Fig. 7B).

The ephaptic intrinsic phase preference 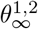 differed substantially between the two cases, converging to (i) 85.1 ms in the reference system and (ii) 185.2 ms when ionic effects were absent. 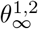 was calculated by taking the mean value over the fifth minute in case (i) and over the fifteenth minute in case (ii), and additionally averaged across all three choices of Δ_lag_. This difference in simulation time was due to the considerably slower convergence of *θ*^1,2^ towards 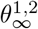 in the absence of ionic ephaptic coupling (the convergence of case (ii) is seen in S4 Fig, showing fifteen minutes of simulation). These differences are not so surprising given that case (i) and (ii) represent quite different systems: “turning off” the ionic ephaptic coupling made neurons independent of the extracellular ion concentration dynamics and reduced the steady-state firing rate from 1.03 Hz to 0.68 Hz.

Our results suggest that the existence of an ephaptic intrinsic phase preference is an electric ephaptic phenomenon, as the results in Fig. 7 show that electric ephaptic effects alone are sufficient to give the system an ephaptic intrinsic phase preference. However, since our framework did not allow us to run simulations with ionic ephaptic effects in isolation (see Section 3.1.3), we cannot exclude the possibility that such a phase preference could arise in the presence of ionic ephaptic effects alone.

#### 2.4.3 The ephaptic intrinsic phase preference follows a trend

To assess whether the existence of an intrinsic ephaptic phase preference 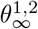 is consistent across different scenarios, we repeated the simulations using a range of stimulus strengths *I*_*c*_ between 16 pA and 40 pA. For the two-neuron system (from Fig. 6 and Fig. 7), we found that the system had an intrinsic ephaptic phase preference 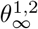 over the entire stimulus range, and that 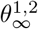 was independent of the initial delay, but decreased with increasing stimulus strength (Fig. 8A). In all cases, the spike phase converged well within five minutes of simulation time (S5 Fig).

**Fig 8.**
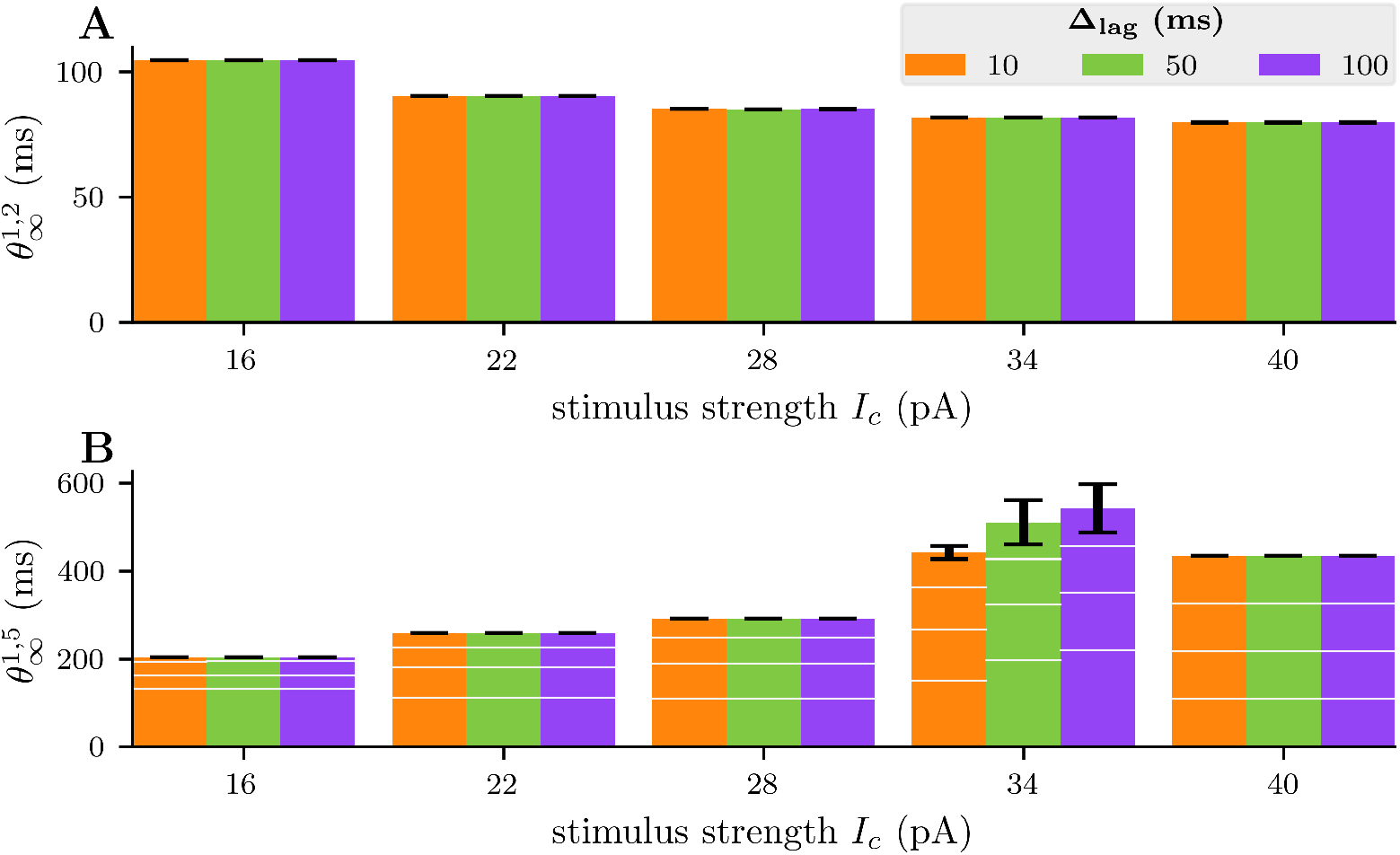
Steady-state spike phases. Each panel shows the converged spike phases 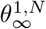, where *N* indicates the number of neurons, using *N* = 2 (**A**) and *N* = 5 (**B**), for a variety of stimulus strengths, *I*_*c*_. The spike phases 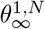 (spike time distance between the first and the *N* ‘th neuron) are calculated as the mean value (standard deviation shown with black bar) over the last minute of the simulation. Each simulation with two neurons in **A**, ran for 5 minutes of simulated time, while the simulations in **B** with five neurons ran for 15 minutes of simulated time, except for the case with *I*_*c*_ = 34 pA. The simulation with *I*_*c*_ = 34 pA ran for 25 minutes of simulated time, without converging (panel D in S6 Fig). White horizontal lines in **B** indicate spike phases 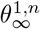 between the first and *n*’th neuron for *n* = 2, 3, 4. All neurons received a constant stimulus, each of strength *I*_*c*_ and turned on with a delay of Δ_lag_ from neuron number *n* to neuron *n* + 1.

We next ran the corresponding simulations for a system of *N* = 5 neurons (Fig. 8B). With five neurons, the system also converged towards an intrinsic ephaptic phase preference 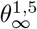 in most cases, with convergence independent of the initial delay. In the five-neuron system, we found that 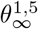 increased with stimulus strength, opposite to the trend observed for 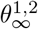 in the two-neuron system. However, the phase difference between the first and second neuron 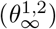 in the five-neuron system decreased with increased stimulus strength, as it did in the two-neuron system. Hence, the increase in 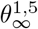 with stimulus strength was due to increased spike phase differences between the subsequent neurons (Fig. 8B).

An exception to the trend was observed for the stimulus *I*_*c*_ = 34 pA. In this case, *θ*^1,5^ did not converge to a stable value in the 25 minutes of simulation. Investigating the simulation (panel D in S6 Fig), we see that *θ*^1,5^ seemed to “almost converge” (around *t* = 10 minutes) to a value that would fit well into the trend observed for the other stimuli in Fig. 8B, but then suddenly started to undergo rather rapid increases. This behavior reflects the inherent complexity of the nonlinear system and indicates that convergence to a fixed point may not occur for all stimulus levels.

We did not examine the exception case further, because our main (and qualitative) conclusion is that ephaptic neuron-ECS systems of the kind studied here can possess ephaptic intrinsic phase preferences which make groups of ephaptically coupled neurons fire in particular sequences that are independent of initial conditions. We have demonstrated the presence of this phenomenon for neural systems of different sizes (*N* = 2 and *N* = 5) and for a range of different stimulus strengths. We have also demonstrated that the phenomenon does not depend critically on the choice of neuron model: we found intrinsic ephaptic phase preferences also when we ran simulations corresponding to those above with the edHH model (see S7 Fig–S10 Fig).

## 3 Discussion and conclusion

In this paper, we have explored the relative roles of electric and ionic ephaptic effects in a small model system consisting of a few neurons interacting through a shared, closed extracellular space. In line with what has been reported in previous studies [9, 31], we found that electric ephaptic coupling did not lead to significant changes in overall firing rates. In contrast, ionic ephaptic effects, mediated by activity-induced changes in extracellular ion concentrations, were shown to have a pronounced impact on firing rates in our model. The influence of extracellular ion concentrations on firing rates and neuronal firing properties has been demonstrated in numerous earlier studies [12, 14–17, 32]. However, these effects have rarely been discussed in the context of ephaptic coupling.

Although ephaptic effects are defined generally as all non-synaptic interactions between neurons [1], some readers may not be familiar with thinking of extracellular ion concentrations as a contributor to ephaptic effects. However, one may argue that the “baseline” extracellular ion concentrations in any brain region represent an equilibrium state reflecting the average level of neuronal (and glial) activity in that region. These “baseline” concentrations would take on different values if, for example, all neurons were silent, and they constitute an ephaptic effect insofar as they feed back onto neuronal dynamics. We may illustrate this with reference to Fig. 3C: When the neurons in our model were inactive, the baseline extracellular K^+^ concentration at the soma depth 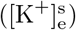 was approximately 6 mM. At an average firing rate of about 2 Hz (green curve), 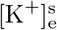 settled to a new steady level of approximately 8 mM. [K^+^]_e_ is tightly regulated, and typically ranges from baseline values and up to 12 mM during intense neuronal activity, while levels above this are associated with pathological conditions [33–35]. The 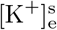 levels in our simulation stayed within this non-pathological range.

While not important for overall firing rates, electric ephaptic effects did cause small changes in spike times of individual neurons. The role of such spike-timing effects in synchronization of neural activity has been the topic of many previous studies [4, 6, 25, 31]. In contrast to conventional synchronization effects, our studies showed that electric ephaptic coupling could give a system of neurons an *ephaptic intrinsic phase preference*, that is, a unique and stable phase difference which was independent of their initial phase differences and depended solely on the input received by the neurons. While stable phase locking due to ephaptic interactions has been reported previously [29, 30], the existence of an ephaptic intrinsic phase preference, emerging from the coupled system itself has, to our knowledge, not been previously described. In principle, the neuronal firing sequences associated with the intrinsic phase preference could serve as a barcode identifier that is unique to the stimulus received by the neurons.

We demonstrated the existence of ephaptic intrinsic phase preference across several model implementations, that is, (i) for systems with either two or five interacting neurons, (ii) for a range of stimulus strengths, and (iii) for two distinct neuron models (edPR and edHH). The phenomenon therefore appears to be general in the sense that it does not depend strongly on specific model choices. We should, however, note that our analysis was limited to highly controlled scenarios involving small numbers of identical neurons sharing a closed extracellular space under constant, noise-free input conditions. It is therefore unclear whether the existence of an ephaptic intrinsic phase preference could be expected under more realistic conditions, or whether it could play a functional role in neural coding.

### 3.1 Model framework

The model framework developed in the current work builds strongly on a model we published previously, comprising a single electrodiffusive Pinsky-Rinzel neuron exchanging ions with a closed extracellular space [17]. We then referred to it as a *minimal model that has it all*. By “has it all”, we in this context meant that it (i) has a spatial extension, (ii) is based on a consistent electrodiffusive framework that accounts for intra- and extracellular dynamics of all ion concentrations and ensures a consistent relationship between these ion concentrations and the electric potential in all compartments, (iii) can have different ion channels in the soma versus dendrites, and (iv) contains the homeostatic machinery that ensures that it maintains a realistic dynamics in ion concentrations during long-time activity.

By “minimal”, we referred to the choice of using only two compartments for the neural extension as well as for the ECS. A single-compartment model would not be able to account for any intracellular or extracellular diffusion nor could it capture extracellular potentials, which arise from the spatial separation of inward and outward transmembrane currents at different membrane locations [2]. The motivation for keeping the number of compartments so low was partly to reduce computational costs and partly to enhance model transparency.

#### 3.1.1 Shared extracellular space

In the current application, we expanded the model presented in Sætra et al. 2020 [17] by including several neurons to study how they interacted ephaptically. Keeping the model “minimal”, we let the neurons share a common (two-compartment) ECS rather than, for example, assigning an individual ECS to each neuron and connecting them via electrodiffusion. For the small group of neurons (ten or fewer) considered in our studies, the assumption that they are in close proximity and experience the same extracellular conditions seems reasonable. However, if this framework is used to model larger neuronal networks, where we cannot assume that changes in one region of the ECS have an immediate effect elsewhere, it would be more appropriate to divide the ECS into multiple interconnected compartments. We demonstrated such a network configuration in Sætra and Mori (2024) [36].

#### 3.1.2 Closed boundary conditions

The closed boundary conditions used for the extracellular space are equivalent to periodic boundary conditions. This means that, for example, when we studied a system of five neurons, it was effectively surrounded by identical “boxes” of five-neuron systems firing at exactly the same times as the five neurons in our box. Only under such conditions would there be no signal going out of the extracellular neighborhood, as there would then be no extracellular gradients between neighboring boxes. Hence, our choice of using closed extracellular boundaries would effectively correspond to a quite high degree of synchrony in our system. Here, we discuss the suitability of using closed boundary conditions for predicting extracellular ion concentrations and electric potentials in our setup.

We do not expect that the closed-boundary condition makes the predicted ion concentration effects unrealistic. Concentrations build up slowly, and rather than depending on exact spike times, they are signatures of the average activity levels in “the neighborhood”. It is not unlikely that neighboring “boxes” in a given brain region should contain neurons with similar activity levels. Open-boundary conditions, where ions were, for example, allowed to leak into an infinite reservoir with fixed concentrations, would probably be a poorer choice for the studies we made here.

Before moving on to discuss electric potentials, we note that in cases of very confined extracellular spaces, such as the narrow gaps between axons in axon bundles, pronounced ion concentration variations - and thus ionic ephaptic effects - can occur on a fast time scale [37]. However, such rapid concentration shifts did not occur at the population-level ECS considered in the current study.

By design, the extracellular potentials produced by neuronal activity in our model system were a function of the extracellular volume. Although we used a realistic ratio between intra- and extracellular volume, the extracellular potential produced by a single action potential was, in our default model implementation, a couple of millivolts. This is much higher than spike amplitudes picked up in extracellular recordings, where spike amplitudes are typically in the order of 0.1 mV [38].

The high spike amplitude observed in our model can, to some degree, be regarded as a boosting effect caused by the effectively periodic boundary conditions in our system. It is, however, uncertain whether this “boosting” means that electric ephaptic effects are overestimated by our model. Extracellular electrodes effectively record the average potential over the surface of the electrode. In contrast, the extracellular spike-amplitude decays as a function of distance from the membrane [2]. How a neuron’s spike will impact its neighboring ephaptically will thus depend on local morphological detail. The impact can, for example, be very strong if the axon or dendrites of one neuron pass very close to the cell body of another [10], and neural tissue is known to be densely packed with average extracellular distances between cellular structures being as low as 10–80 nm [39, 40].

Morphological details like this are not captured by the coarse model setup used in our study. However, we do not necessarily believe that the unrealistically large extracellular spike amplitudes in our model imply that the electrical ephaptic effects it predicts are unrealistic.

#### 3.1.3 Isolating ionic ephaptic effects

The KNP formalism ensures a self-consistent relationship between the ionic concentrations, the associated electric charge, and the electric potential in all model compartments. Because the equations governing the electric potential in the different model compartments are coupled (see Section 4.2.2), we were not able to “turn off” electric ephaptic effects without compromising the consistent computation of the intracellular potentials and membrane potentials.

A limitation of using this self-consistent framework is that ionic ephaptic effects could not be studied in isolation. In contrast, electric ephaptic effects could be isolated, as ionic ephaptic effects could be eliminated by making membrane mechanisms independent of ion-concentration variation by instead fixing reversal potentials and pump rates to constants derived from baseline concentrations rather than allowing them to vary with the concentration variables.

Given these constraints, the roles of ionic ephaptic effects were assessed indirectly by comparing scenarios with (i) both ionic and electric ephaptic effects present, (ii) only electric ephaptic effects present, and (iii) effectively no ephaptic effects present, as achieved by letting the ECS be very large. This approach allowed us, for example, to conclude that ionic (but not electric) ephaptic effects were important for regulating system firing rates, as summarized in Fig. 4.

However, the need to assess the role of ionic ephaptic effects indirectly did have some limitations. For example, while a comparison of the scenarios (i)–(iii) in Fig. 7 demonstrated that electric ephaptic effects are sufficient for obtaining an ephaptic intrinsic phase preference, it did not allow us to exclude the possibility that such a phase preference could arise in the presence of ionic ephaptic effects alone.

### 3.2 Outlook

A very common model simplification used in computational neuroscience is that ion concentrations remain constant during the simulated period. Making this assumption, the effects of concentration variations on ionic reversal potentials, or of ionic diffusion on electrical potentials, are not accounted for. In many cases, this may not be critical for the model’s predictions, since ion concentrations tend to vary quite little during normal working conditions.

We have, for more than a decade, developed ion-conserving computational models for conditions where the constant-concentration assumption does not hold [17–20, 36, 37, 41–46], drawing inspiration from previous works on electrodiffusion in brain tissue [47–49]. Electrodiffusive models of this kind are computationally expensive, and many important questions in neuroscience are probably addressed more wisely using more conventional models based on e.g., the NEURON simulation environment [50, 51].

However, ion-conserving models are needed for studying conditions in the brain in which extracellular ion concentrations are expected to vary significantly in time and space. While such variations do occur during non-pathological periods of intense neural signaling [11], they may play a particularly important role for brain functioning under pathological conditions. Dramatic shifts in extracellular ion concentrations are a hallmark for conditions such as spreading depression, epilepsy, and anoxia [11, 52–54]. Notably, epilepsy (and to some degree spreading depression) is also associated with a large degree of synchrony among the involved (or affected) neurons. Since these pathologies are not highly localized, but involve similar activity in rather extended brain regions, the (in practice) periodic-boundary conditions used in the current study may be a plausible choice if one were to address such pathologies in a modeling study. In conclusion, we do believe the model presented in the current work, and other models of this kind, will be important for studying brain conditions associated with changes in extracellular ion concentrations.

## 4 Methods

The network model used in this paper is based on the edPR model presented in Sætra et al. (2020) [17], a multicompartmental neuron model with homeostatic mechanisms and electrodiffusive ion concentration dynamics. While the original edPR model includes a single cell, we here extend the model to include an arbitrary number of neurons embedded in a shared ECS (Fig. 1). Each neuron is described by two compartments, one representing the soma and one representing the dendritic tree. The neurons are indirectly connected through a shared ECS, which is also divided into two compartments. All neurons are oriented in the same direction, such that the *upper* ECS compartment is in direct contact with the dendritic compartment of all neurons, while the *lower* ECS compartment is in direct contact with all somatic compartments (Fig. 1). Via a set of coupled ordinary differential equations (ODEs), the model predicts the temporal evolution of the electric potential and ion concentrations in each compartment.

### 4.1 Notation

We will be using the following notation: An unspecified ion species is represented by *k*, with its concentration denoted [*k*]. A function or variable *f* specific to ion species *k* is marked as *f*_*k*_. Similarly, *f* is specified by subscripts “i” or “e” to belong to the intracellular space, as *f*_i_, or to the extracellular space, as *f*_e_. Both ion species and region may be specified simultaneously, using *f*_*k*,e_ and *f*_*k*,i_. Further, superscripts “s” and “d” indicate the soma or dendrite layer, respectively. This means that 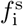 corresponds to a soma compartment, while 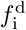 corresponds to a dendrite compartment. Note that the two extracellular compartments are also marked with superscripts “s” and “d”, to identify the ECS compartment in direct contact with the somas, 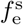, and the ECS compartment in direct contact with the dendrites, 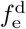. Lastly, individual neurons are labeled with *n* = 1, 2, …, *N*, where *N* is the number of neurons. This way, 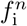 is a neuron-specific function or variable. To avoid clutter, the neuron index *n* is omitted when not explicitly needed.

### 4.2 The Kirchhoff-Nernst-Planck framework

To predict ion concentrations and electric potentials, the model uses the KNP framework, ensuring a biophysically consistent relationship between ion concentrations, electric charge, and electric potentials [17, 19, 41].

#### 4.2.1 Electrodiffusion

The flux of ion species *k* from the soma layer to the dendrite layer is divided into two components:

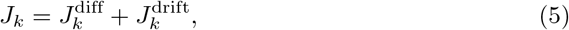

describing electrodiffusion. While *J*_*k*,e_ is the flux between the extracellular compartments, there are *N* intracellular fluxes *J*_*k*,i_, one for each neuron.

Firstly, the diffusive component depends on the ion concentrations [*k*], as given by

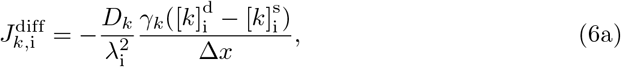

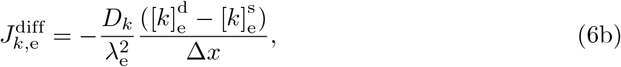

where *D*_*k*_ is the diffusion constant, *γ*_*k*_ is the fraction of free ions in the ICS, *λ*_i_ and *λ*_e_ are the tortuosities, and Δ*x* is the intercompartmental distance.

Secondly, the electric-drift component additionally depends on the electric potentials *ϕ*, according to

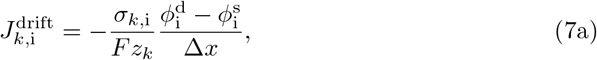

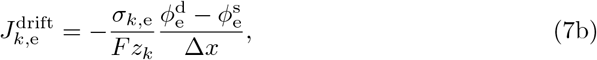

where *σ*_*k*,i_ and *σ*_*k*,e_ are the ion-specific conductances, *F* is the Faraday constant, *R* is the universal gas constant, *T* is the absolute temperature, and *z*_*k*_ is the signed valency of ion species *k*. The conductances are given by

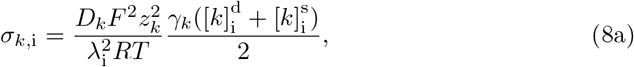

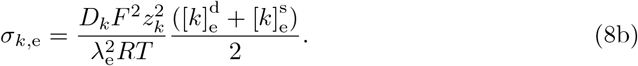

#### 4.2.2 Electric potentials

The membrane potentials of each neuron, at the dendrite 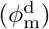 and soma 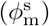, are defined in accordance with the charge-capacitor relationship. This means that the membrane potentials are given as the total charge divided by the total capacitance, i.e.,

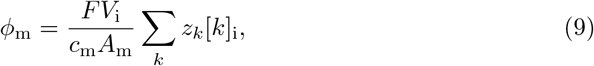

where *c*_m_ is the capacitance per area, *A*_m_ is the membrane area of a single neuron compartment, and *V*_i_ is the neuron compartment volume.

The extracellular compartment at dendritic depth is chosen as the reference point for the electric potential, such that

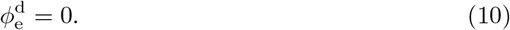

Further, the electric potential of the extracellular compartment at the soma depth is found by requiring that currents travel in closed loops. This implies that the net extracellular electric current (up or down) must be equal in magnitude and opposite in direction to the net intracellular electric current summed over all neurons *n*:

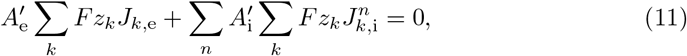

where 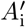 and 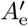 are the intracellular and extracellular cross-sections areas, respectively. From this, it follows that

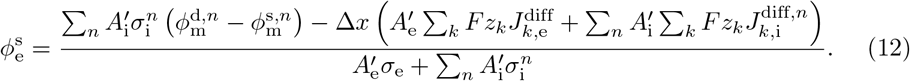

Lastly, by using the definition *ϕ*_m_ = *ϕ*_i_ − *ϕ*_e_, we get the expressions for the intracellular electric potentials as

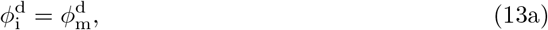

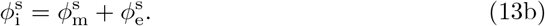

For a more detailed derivation of the electric potentials (of the single-neuron case), we refer the reader to Sætra et al. (2020) [17].

#### 4.2.3 Ion concentrations

The time evolution of the ion concentrations [*k*] assumes conservation of ions. All compartment concentrations are calculated for each of the ion species except for X^−^, which is an immobile ion introduced to ensure electroneutrality.

The intracellular concentrations of a neuron evolve according to

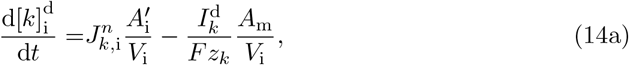

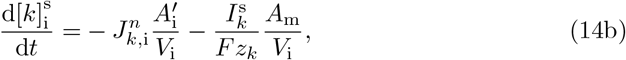

where 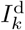 and 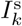 are the transmembrane currents, which will be addressed in the next section.

Similarly, the time evolution of the ion concentrations in the extracellular compartments is determined by

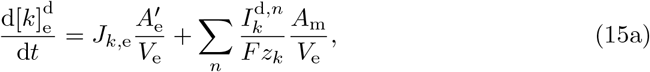

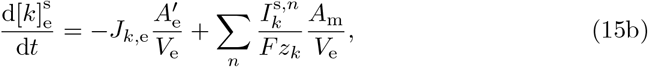

where *V*_e_ is the extracellular compartment volume, and the sum goes over all neurons in the system.

The equations are solved numerically, requiring initial conditions for all ion species in every compartment. Unless otherwise stated, the results produced in this paper used steady-state values for the initial conditions (provided in Section 4.6).

### 4.3 Transmembrane currents

Active and passive ion channels follow the general formula:

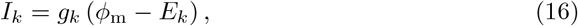

where *g*_*k*_ is the conductance, with a maximum conductance 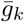, and *E*_*k*_ is the reversal potential given by

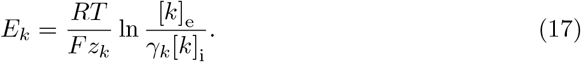

The passive, ion-specific leak channels are given by

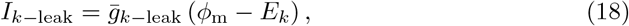

for Na^+^, K^+^, and Cl^−^ .

Specific active ion channels are described in later sections (as well as in S1 Text for the edHH model), though their gating variables often evolve according to the rate law of first-order kinetics. The general equation for the time evolution of a gating variable *ν* is given by

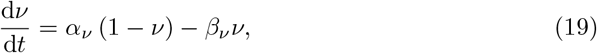

where *α*_*ν*_ and *β*_*ν*_ are the rate coefficients of opening and closing the gating particle.

In addition, each neuron may experience a stimulus current *I*_stim_, which we model as a pure K^+^ transmembrane current into the soma. By letting the stimulus current be transmembrane, we ensure that no charge enters or leaves the system [17]. The exact form of *I*_stim_ may vary, and is described for each study in the results section.

Because the purpose of this study is to investigate ephaptic coupling, the neurons will not share any synaptic connections but rather be *indirectly* connected through the extracellular space. For future investigations, synapses can easily be added to the model, see e.g. Sætra et al. (2024) [36].

#### 4.3.1 Homeostatic mechanisms

In addition to the leak- and active ion channels, the model incorporates homeostatic mechanisms. Their functionals and parameters are equivalent to those of the single-neuron edPR model presented in Sætra et al. (2020) [17].

First we have the 3Na^+^/2K^+^-pump,

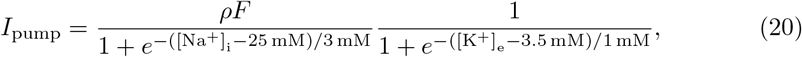

where *ρ* is the pump strength. Next, we have two co-transporters: the K^+^/Cl^−^ (KCC2) co-transporter, given by

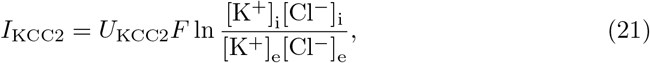

and the Na^+^/K^+^/2Cl^−^ (KNCC1) co-transporter, given by

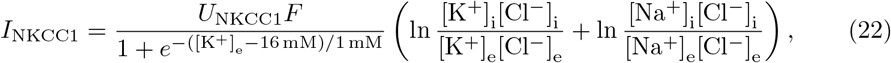

where *U*_KCC2_ and *U*_NKCC1_ are the co-transporter strengths. Lastly, we have intracellular Ca^2+^ decay (Ca-dec). The Ca^2+^ decay was modeled using

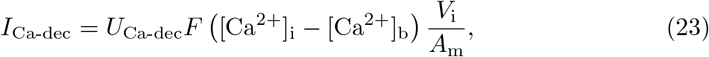

where *U*_Ca-dec_ is the decay rate and [Ca^2+^]_b_ is the intracellular basal concentration of calcium ions.

#### 4.3.2 Dendrite membrane model

The edPR model contains active ion channels at both the dendrites and the soma, in addition to all homeostatic mechanisms, leak channels, and the stimulus channel, as described above.

Three active channels are included for the dendrite membranes: two K^+^ channels and one calcium channel. First, we have the potassium after-hyperpolarisation (K-AHP) channel, given by

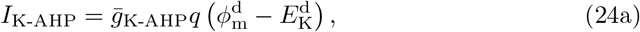

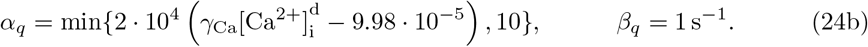

where 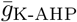 is the maximum conductance, and *q* is a gating variable with calcium-dependent rate coefficients, evolving according to the rate law in Eq. (19). The second potassium channel is calcium-gated (K-C):

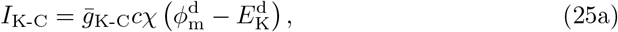

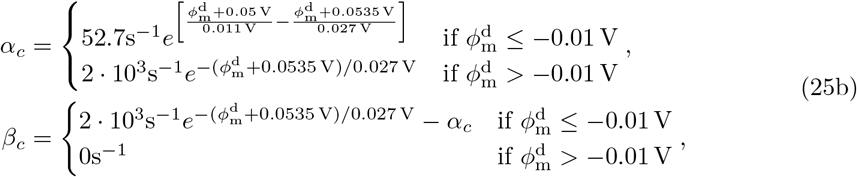

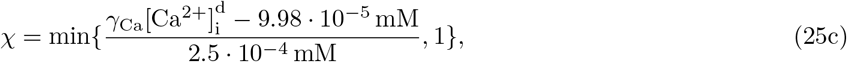

with maximum conductance 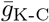, and where *c* is a gating variable following Eq. (19), and *χ* introduces the calcium-dependency of the channel. Lastly, there is the voltage-dependent calcium channel given by

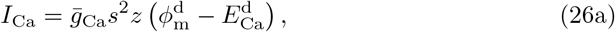

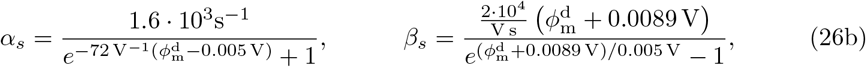

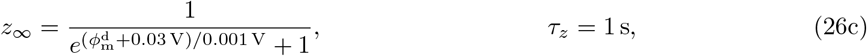

with maximum conductance 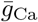, and where gating variable *s* follows the rate law in Eq. (19), while gating variable *z* evolves according to d*z/*d*t* = (*z*_∞_ − *z*)*/τ*_*z*_. The overview of the dendritic edPR membrane mechanisms is then

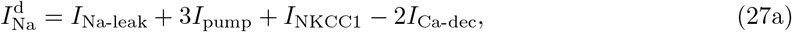

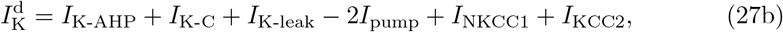

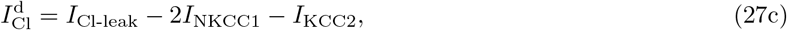

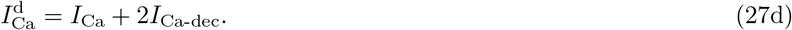

#### 4.3.3 Soma membrane model

At the soma, there are two active channels in the edPR model. First, an active sodium channel, given by

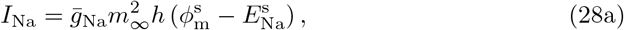

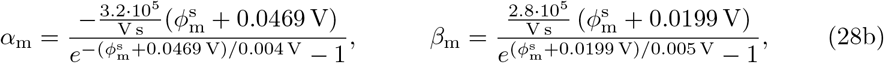

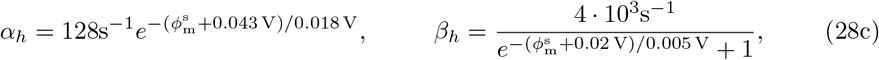

where 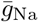 is the maximum conductance, *m*_∞_ is the steady-state gating variable given by *m*_∞_ = *α*_m_*/*(*α*_m_ + *β*_m_), and *h* evolves according to the rate law given in Eq. (19). Secondly, there is an active potassium channel, the delayed rectifier (K-DR):

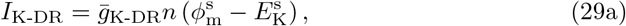

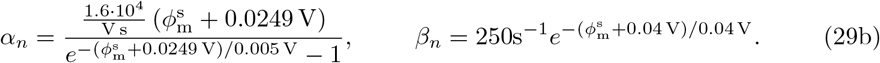

Here, 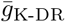 is the maximum conductance and *n* evolves according to Eq. (19). The overview of edPR membrane channels at the soma is provided below.

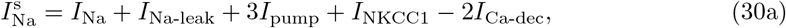

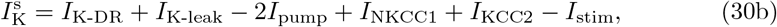

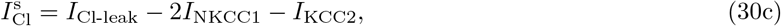

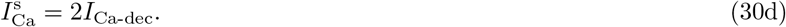

### 4.4 Model parameters

Ion-specific parameters are given in Table 1. Geometrical parameters and temperature are given in Table 2. Lastly, the membrane parameters are listed in Table 3.

**Table 1.** Diffusion constants, fraction of free intracellular ions, and valency. Values are taken from the original single-neuron model [17].

| $\mathbf{k}$ | $\mathbf{D_k}$ | $\gamma_{\mathbf{k}}$ | $\mathbf{z_k}$ |
| --- | --- | --- | --- |
| $\text{Na}^+$ | $1.33 \cdot 10^{-9} \text{ m}^2 \text{ s}^{-1}$ | 1 | 1 |
| $\text{K}^+$ | $1.96 \cdot 10^{-9} \text{ m}^2 \text{ s}^{-1}$ | 1 | 1 |
| $\text{Cl}^-$ | $2.03 \cdot 10^{-9} \text{ m}^2 \text{ s}^{-1}$ | 1 | -1 |
| $\text{Ca}^{2+}$ | $0.71 \cdot 10^{-9} \text{ m}^2 \text{ s}^{-1}$ | 0.01 | 2 |
| $\text{X}^-$ | $0 \text{ m}^2 \text{ s}^{-1}$ | 1 | -1 |

**Table 2.** Geometry, tortuosities and temperature. All values are taken from [17]. The ECS size scales with the number of neurons in the system, *N* .

|  | Value | Description |
| --- | --- | --- |
| $\Delta x$ | $6.67 \cdot 10^{-4} \text{ m}$ | Soma-to-dendrite distance (ICS and ECS) |
| $A_m$ | $6.16 \cdot 10^{-10} \text{ m}^2$ | Compartment membrane area |
| $A'_i$ | $12.32 \cdot 10^{-10} \text{ m}^2$ | Cross-section area between soma and dendrite |
| $A'_e$ | $N/2 \cdot A'_i$ | Cross-section area between ECS compartments |
| $V_i$ | $1.437 \cdot 10^{-15} \text{ m}^3$ | Intracellular compartment volume |
| $V_e$ | $N/2 \cdot V_i$ | Extracellular compartment volume |
| $\lambda_i$ | 3.2 | Intracellular tortuosity |
| $\lambda_e$ | 1.6 | Extracellular tortuosity |
| $T$ | 309.14 K | Absolute temperature |

**Table 3.** Membrane parameters. All conductances are taken from [17].

|  | Value | Description |
| --- | --- | --- |
| $c_m$ | $0.03 \text{ F m}^{-2}$ | Capacitance per area |
| $\bar{g}_{\text{Na-leak}}$ | $0.247 \text{ Sm}^{-2}$ | Leak conductance |
| $\bar{g}_{\text{K-leak}}$ | $0.5 \text{ Sm}^{-2}$ | Leak conductance |
| $\bar{g}_{\text{Cl-leak}}$ | $1.0 \text{ Sm}^{-2}$ | Leak conductance |
| $\rho$ | $1.87 \cdot 10^{-6} \text{ mol m}^{-2} \text{ s}^{-1}$ | Pump strength |
| $U_{\text{KCC2}}$ | $7.0 \cdot 10^{-7} \text{ mol m}^{-2} \text{ s}^{-1}$ | Co-transporter strength |
| $U_{\text{NKCC1}}$ | $2.33 \cdot 10^{-7} \text{ mol m}^{-2} \text{ s}^{-1}$ | Co-transporter strength |
| $U_{\text{Ca-dec}}$ | $75 \text{ s}^{-1}$ | Calcium decay rate |
| $[\text{Ca}^{2+}]_b$ | $0.01 \text{ mM}$ | Basal intracellular $\text{Ca}^{2+}$ concentration |
| $\bar{g}_{\text{Na}}$ | $300 \text{ Sm}^{-2}$ | Maximum conductance |
| $\bar{g}_{\text{K-DR}}$ | $150 \text{ Sm}^{-2}$ | Maximum conductance |
| $\bar{g}_{\text{K-AHP}}$ | $8 \text{ Sm}^{-2}$ | Maximum conductance |
| $\bar{g}_{\text{K-C}}$ | $150 \text{ Sm}^{-2}$ | Maximum conductance |
| $\bar{g}_{\text{Ca}}$ | $118 \text{ Sm}^{-2}$ | Maximum conductance |

### 4.5 Controlling the ephaptic effects

If we are to determine how ephaptic coupling *changes* the behavior of neurons, then we need to somehow control the factors that lead to ephaptic coupling. This section describes how our framework was manipulated to eliminate ephaptic effects.

#### 4.5.1 Weakening all ephaptic effects

It is safe to assume that the distance between neurons affects the magnitude of ephaptic effects; the closer the neurons are, the more they experience local changes in the environment due to neighboring neurons’ activity. In our KNP framework, there is no explicit distance between neurons, but rather an implicit one determined by the ECS size. Changes in ion concentrations and electric potentials in the extracellular space are instantly homogeneously distributed inside each of its two compartments. Therefore, each neuron will instantly experience the exact same changes in the environment, regardless of which neuron caused the deviance in electric potentials and/or ionic concentrations in the ECS. In the limit of an infinitely large extracellular space, all ephaptic effects, including self-ephaptic effects, are effectively removed.

Such expansion of the ECS is achieved by a factor multiplication of the ECS compartment volumes *V*_e_ and the cross-sectional area 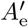 between the two ECS compartments. With a sufficiently large increase, all ephaptic effects are eliminated.

#### 4.5.2 Eliminating ionic ephaptic effects

Electric ephaptic coupling arises from perturbations in the extracellular electric field, directly impacting the membrane potential. Ionic ephaptic coupling is caused by deviations in the extracellular ionic concentrations, which, in turn, affect membrane currents. Pumps, co-transporters, and ion-specific channels are concentration-dependent functionals. The ion-specific channels typically depend on the concentrations through the reversal potential.

To eliminate the ionic ephaptic coupling, the dynamic extracellular ion concentrations in all transmembrane currents are replaced with constant values, fixed at the system equilibrium values. The only exception is Ca^2+^, which remains dynamic in the expressions for the transmembrane currents.

Ions move in and out of the extracellular compartments without changes in the extracellular concentration impacting the membrane currents. The currents are still affected by the evolution of intracellular concentrations, though we did not consider this to be an ephaptic effect. This way, the neurons are prevented from experiencing any changes to the ion concentrations in the ECS. In effect, the ionic ephaptic coupling is removed from the system, leaving only electric ephaptic effects.

### 4.6 Computational details

Both model versions were implemented in the Python programming language, version 3.10.4. The discrete integration method used to solve the coupled ODEs was a 4th-order Runge-Kutta method, using the 3/8-rule. All numerical experiments were carried out with a time step of 2 · 10^−5^ s, while the results were sampled every 10^−4^ s. See S11 Fig and S12 Fig for verification of the time step convergence.

The physical constants *R* and *F* were obtained from the scipy library, and the stochasticity in the noisy stimulus current was obtained using the numpy library. For full implementation, see https://github.com/eirillsh/electrodiffusive-neurons.

Initial values in the simulations were set to steady-state values obtained by a process described in S3 Text. A list of the steady-state ion concentrations, membrane potentials, and gating variables is found in Table 4. In addition, each simulation was run for five seconds without any stimulus current. These five seconds were discarded.

**Table 4.** Steady-state values.

| | $[\mathbf{k}]_i^s$<br>(mM) | $[\mathbf{k}]_e^s$<br>(mM) | $[\mathbf{k}]_i^d$<br>(mM) | $[\mathbf{k}]_e^d$<br>(mM) |
| --- | --- | --- | --- | --- |
| $\text{Na}^+$ | 16.9 | 141 | 16.9 | 141 |
| $\text{K}^+$ | 140 | 5.94 | 140 | 5.94 |
| $\text{Cl}^-$ | 5.43 | 107 | 5.43 | 107 |
| $\text{Ca}^{2+}$ | 0.0100 | 1.10 | 0.0100 | 1.10 |
|  | **soma** |  | **dendrite** |  |
| $\phi_m$ | -67.7 mV | | -67.7 mV | |
| $n$ | 0.000262 | | - | |
| $h$ | 0.999 | | - | |
| $s$ | - | | 0.00716 | |
| $c$ | - | | 0.00527 | |
| $q$ | - | | 0.0107 | |
| $z$ | - | | 1.00 | |

## 5 Supporting information

**S1 Text. Hodgkin-Huxley membrane model**. We present the edHH model, with membrane mechanisms and model parameters.

**S2 Text. Optimal ISI length**. A discussion linking the timing of ephaptic interactions to the ephaptic intrinsic phase preference.

**S3 Text. Steady-state calibration**. We present the steady-state calibration process of both the edHH and edPR models.

## 6 Acknowledgments

The authors would like to thank Marie E. Rognes for insightful discussions during the initial phase of the project. G. T. E. received funding from the European Union’s Research and Innovation Program Horizon Europe under Grant Agreement No. 101147319 [EBRAINS 2.0].

## 7 Author contributions

**Conceptualization:** E. Hauge, G. T. Einevoll, and G. Halnes.

**Formal analysis:** E. Hauge.

**Investigation:** E. Hauge and G. Halnes.

**Methodology:** E. Hauge, M. J. Sætra, and G. Halnes

**Software:** E. Hauge.

**Supervision:** M. J. Sætra, G. T. Einevoll, and G. Halnes.

**Validation:** E. Hauge.

**Visualization:** E. Hauge.

**Writing – original draft:** E. Hauge and G. Halnes.

**Writing – review & editing:** E. Hauge, M. J. Sætra, G. T. Einevoll, and G. Halnes.

## Supporting Figures

**S1 Fig.**
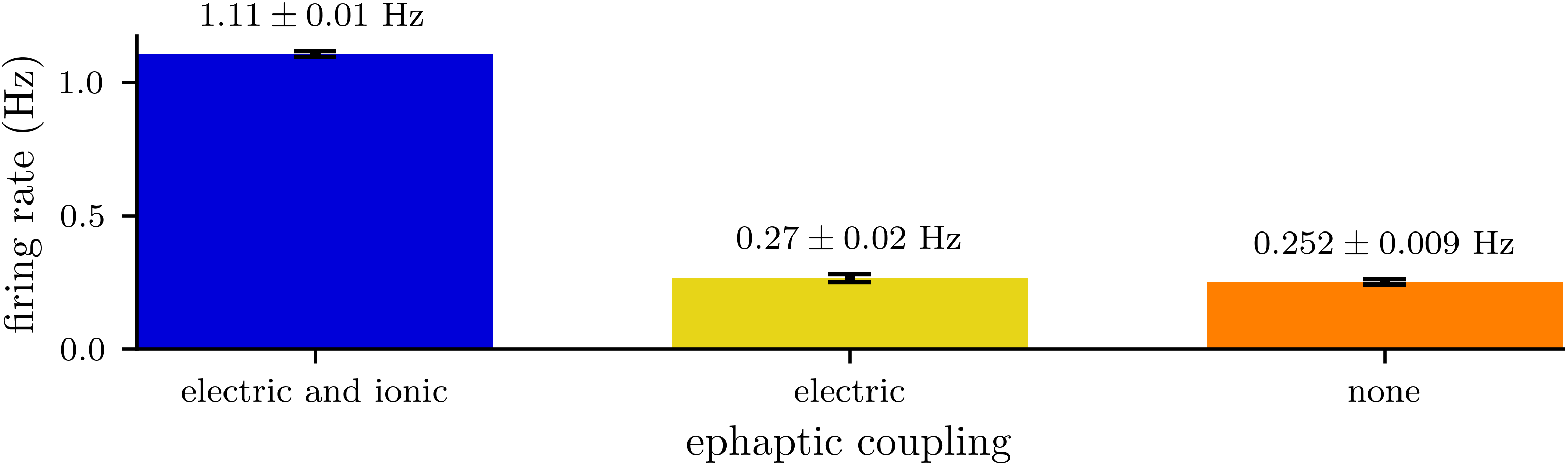
Population firing rates for ten edHH neurons with and without ephaptic coupling. A comparison of firing rates between systems with ephaptic coupling (blue), with electric but without ionic ephaptic coupling (yellow), and without any ephaptic coupling (orange). Firing rates were calculated as the population average over the fifth minute in a system of ten neurons with an average noisy stimulus of 3.75 pA. Removing all ephaptic effects substantially lowered the firing rate, from a reference rate of 1.11 ±0.01 Hz to 0.252 ±0.009 Hz. In accordance with results using the edPR model (Fig. 4), the firing rate of the edHH model with only electric ephaptic coupling (0.27 ±0.02 Hz) was similar to when all ephaptic coupling was removed from the system.

**S2 Fig.**
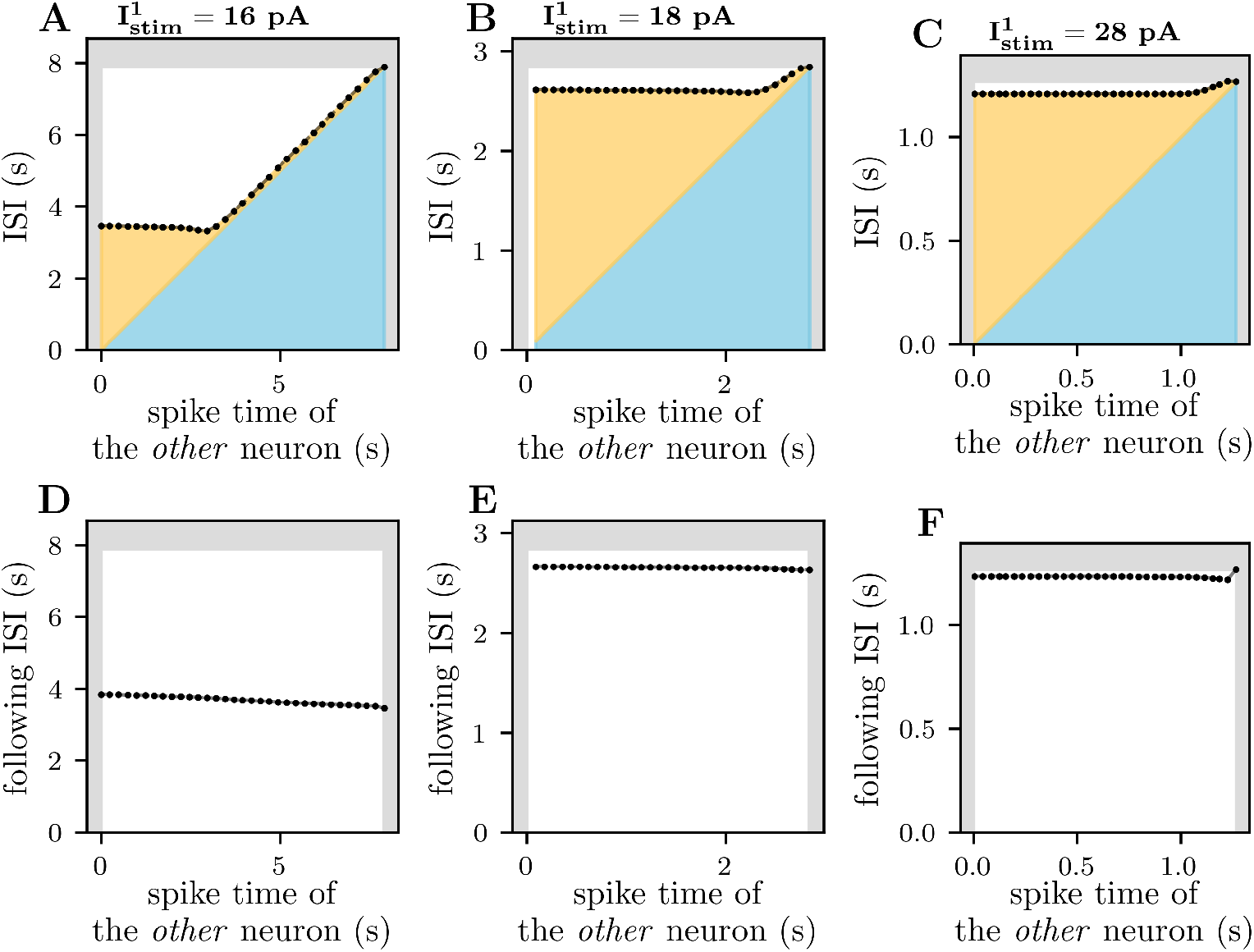
Current and following ISI of an edPR neuron versus the timing of an ephaptic kick. A figure illustrating the impact of the timing of a single ephaptic kick on the current ISI (**A** – **C**) and the following ISI (**D** – **E**). Each panel shows the ISI of the first neuron, either current ISI 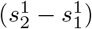 or the following ISI 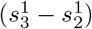, plotted as a function of the second neuron’s spike time 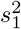, where 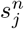 is the *j*’th spike time of the *n*’th neuron. Gray shading illustrates the reference ISI, where the second neuron does not fire. Each column represents a different stimulus strength on the first neuron, with 16 pA corresponding to 0.11 Hz on first column (**A** and **D**), 18 pA correpsonding to 0.35 Hz on the second (**B** and **E**), and 28 pA corresponding to 0.79 Hz (**C** and **F**) on the third column. In panels **A**–**C**, the ISI of the first neuron is shaded light blue and yellow, corresponding to the intervals before and after the ephaptic kick (second neuron spikes), respectively. The impact of the spike time of the *other* neuron, 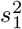, on the ISI 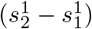 decreased with stronger stimulus current on the first neuron (**A**–**C**). This means that, at higher firing rates, the timing of the ephaptic coupling is less important compared to neurons with lower firing rates. While the timing of the ephaptic kick impacted the *current* ISI, the *following* ISI was barely impacted by the exact time of 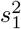 (**D**–**F**). The succeeding ISI was shortened by ephaptic coupling, but the exact timing was less important.

**S3 Fig.**
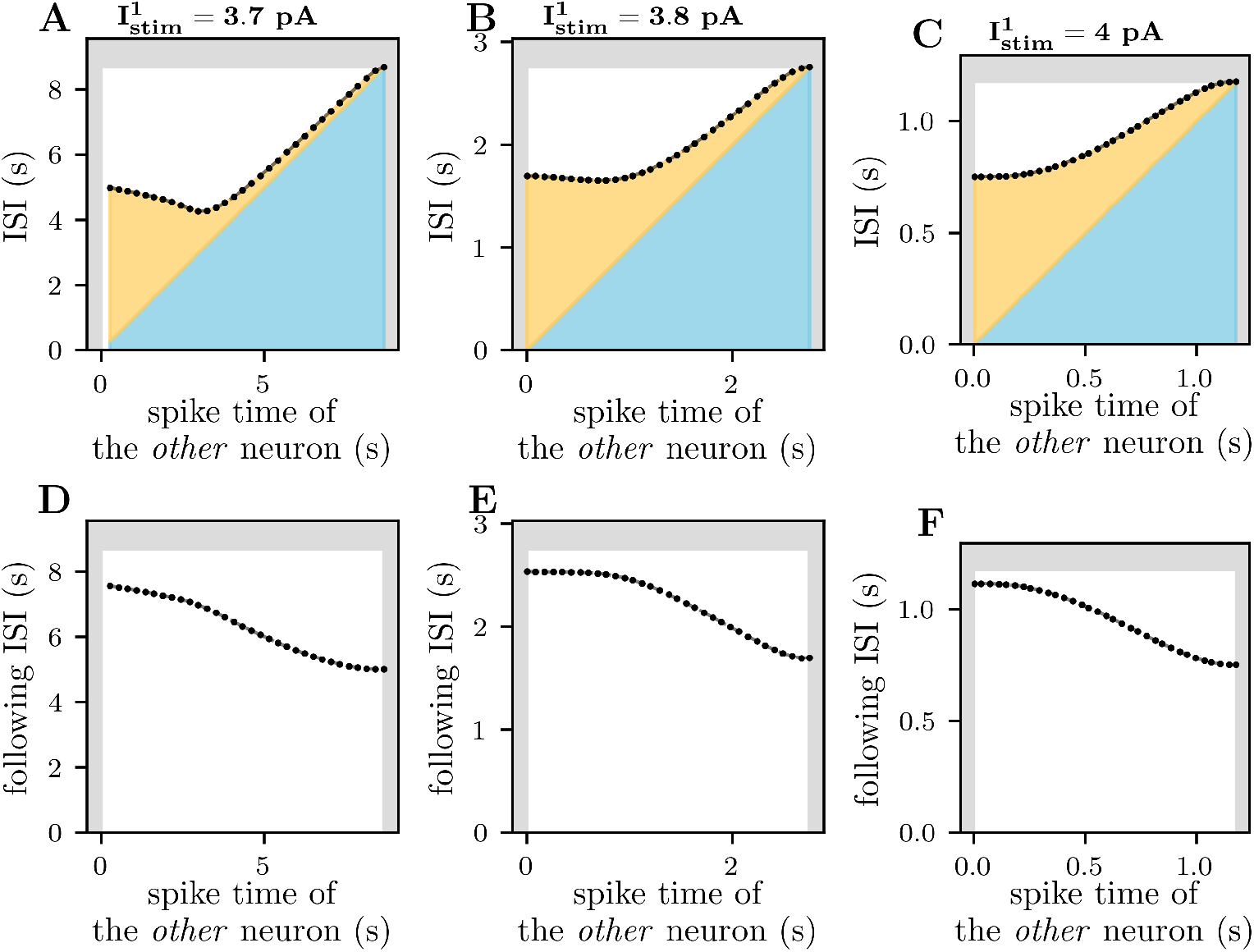
Current and following ISI of an edHH neuron versus the timing of an ephaptic kick. A figure illustrating the impact of the timing of a single ephaptic kick on the current ISI (**A** – **C**) and the following ISI (**D** – **F**). Each panel shows the ISI of the first neuron, either current ISI 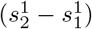 or the following ISI 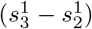, plotted as a function of the second neuron’s spike time 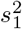, where 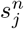 is the *j*’th spike time of the *n*’th neuron. Gray shading illustrates the reference ISI, where the second neuron did not fire. In panels **A**–**C**, the ISI of the first neuron is shaded light blue and yellow, corresponding to the intervals before and after the ephaptic occurrence (second neuron spikes), respectively. Each column represents a different stimulus strength on the first neuron, with 3.7 pA corresponding to a firing rate of 0.12 Hz in the first column (**A** and **D**), 3.8 pA corresponding to 0.37 Hz in the second (**B** and **E**), and 4.0 pA corresponding to 0.85 Hz in the third column (**C** and **F**). In agreement with the results obtained using the edPR model (S2 Fig), the exact spike time of the other neuron had a larger impact on the ISI at lower firing rates (**A**–**C**). However, the impact of the timing was, in general, greater when using the edHH model compared to the simulations using the edPR model. In contrast to simulations using edPR, the timing of 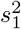 also impacted the following ISI (**D**–**F**) when using the edHH model.

**S4 Fig.**
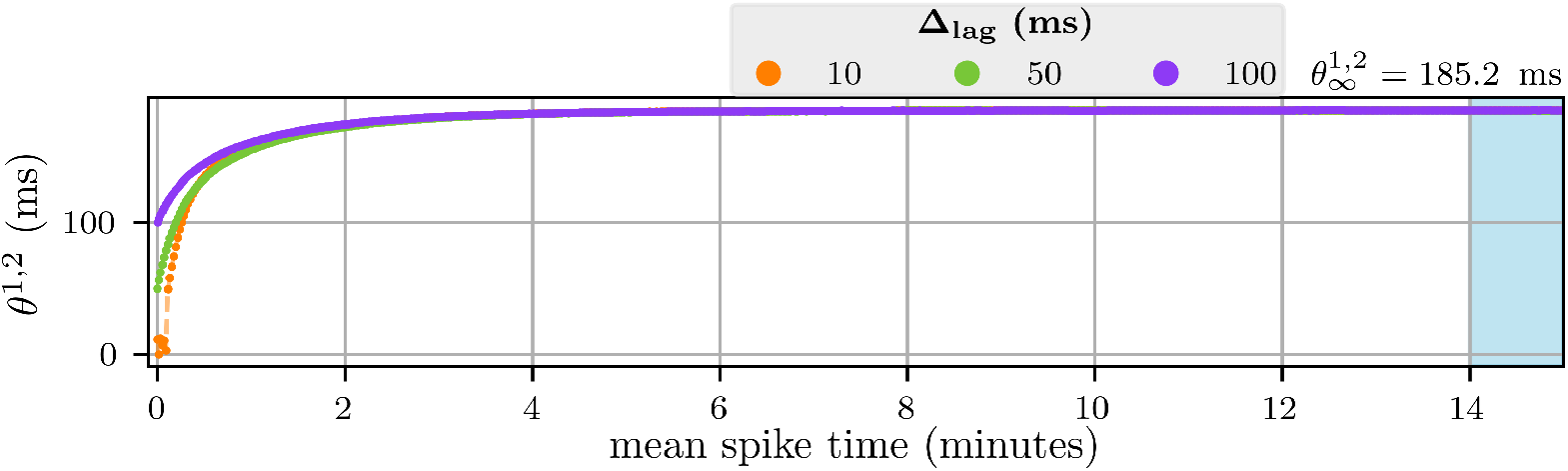
The spike phase difference of two edPR neurons in the absence of ionic ephaptic coupling. A figure displaying the time evolution of the phase difference 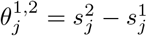 of two edPR neurons, where 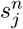 is the *j*’th spike time of the *n*’th neuron. Both neurons received an equally strong constant stimulus current of 28 pA, which the first neuron received from the beginning of the simulation, while the onset of the second neuron’s stimulus started at time Δ_lag_. Among all three cases of Δ_lag_, the population firing rate was 0.68 Hz. Both the converged spike phase 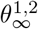 and the firing rates were calculated as the mean over the fifteenth minute (shaded blue area). The neurons were prevented from experiencing changes in the extracellular ion concentrations.

**S5 Fig.**
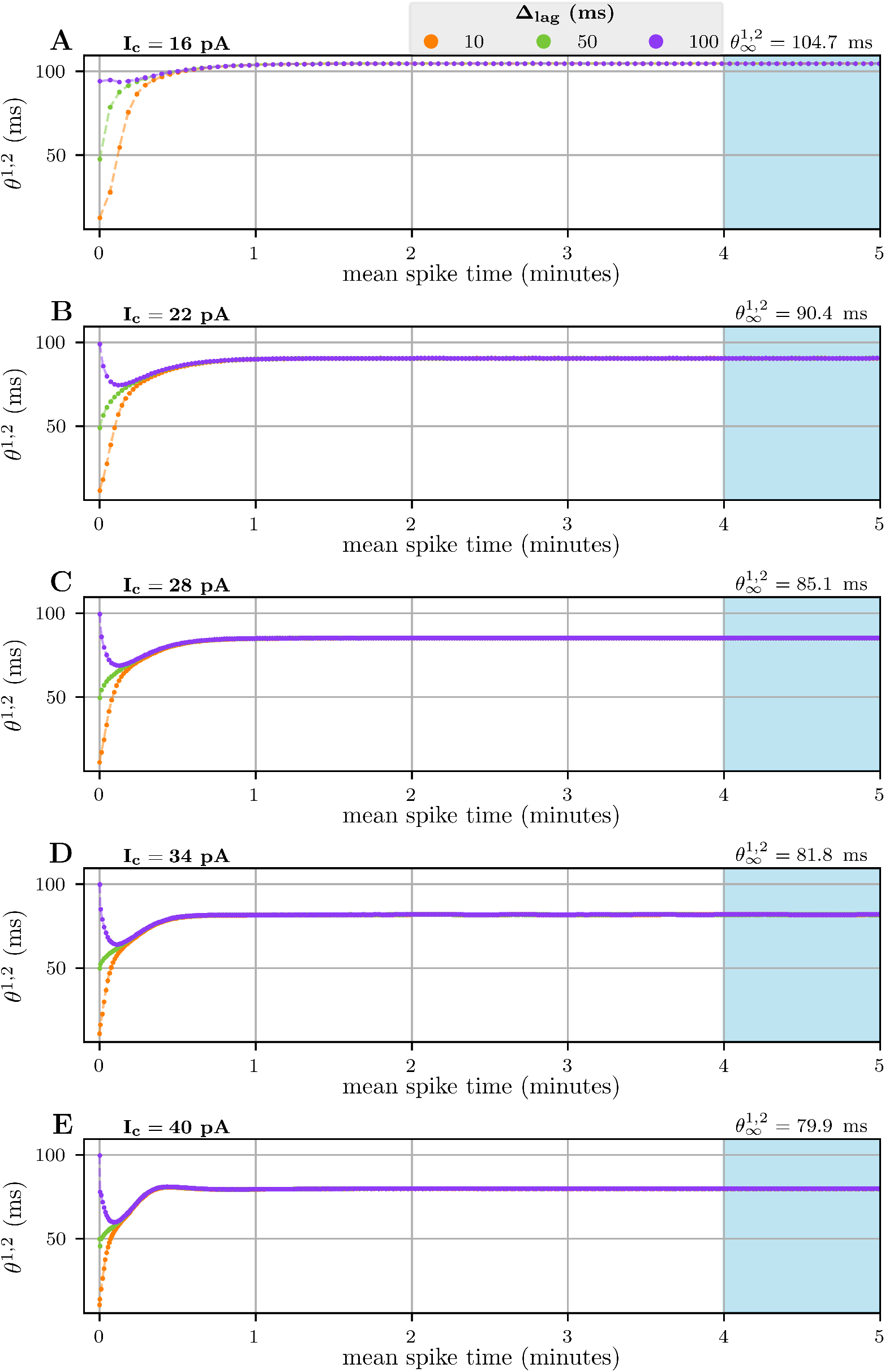
The spike phase difference of two edPR neurons over time. A figure displaying the phase difference *θ*^1,2^ as a function of time. Each panel (**A**–**E**) plots the time evolution of 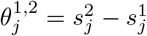, where 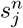 is the *j*’th spike time of the *n*’th neuron. Both neurons received an equally strong constant stimulus current with strength *I*_*c*_, which gave a population firing rate of 0.33 Hz in **A**, 0.70 Hz in **B**, 1.03 Hz in **C**, 1.38 Hz in **D**, and 1.78 Hz in **E**, which were all independent of the chosen initial delay Δ_lag_. The first neuron received the stimulus from the beginning of the simulation, while the onset of the second neuron’s stimulus was delayed by Δ_lag_. Converged spike phases 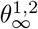 and firing rates were calculated as the mean over the fifth minute (shaded blue area).

**S6 Fig.**
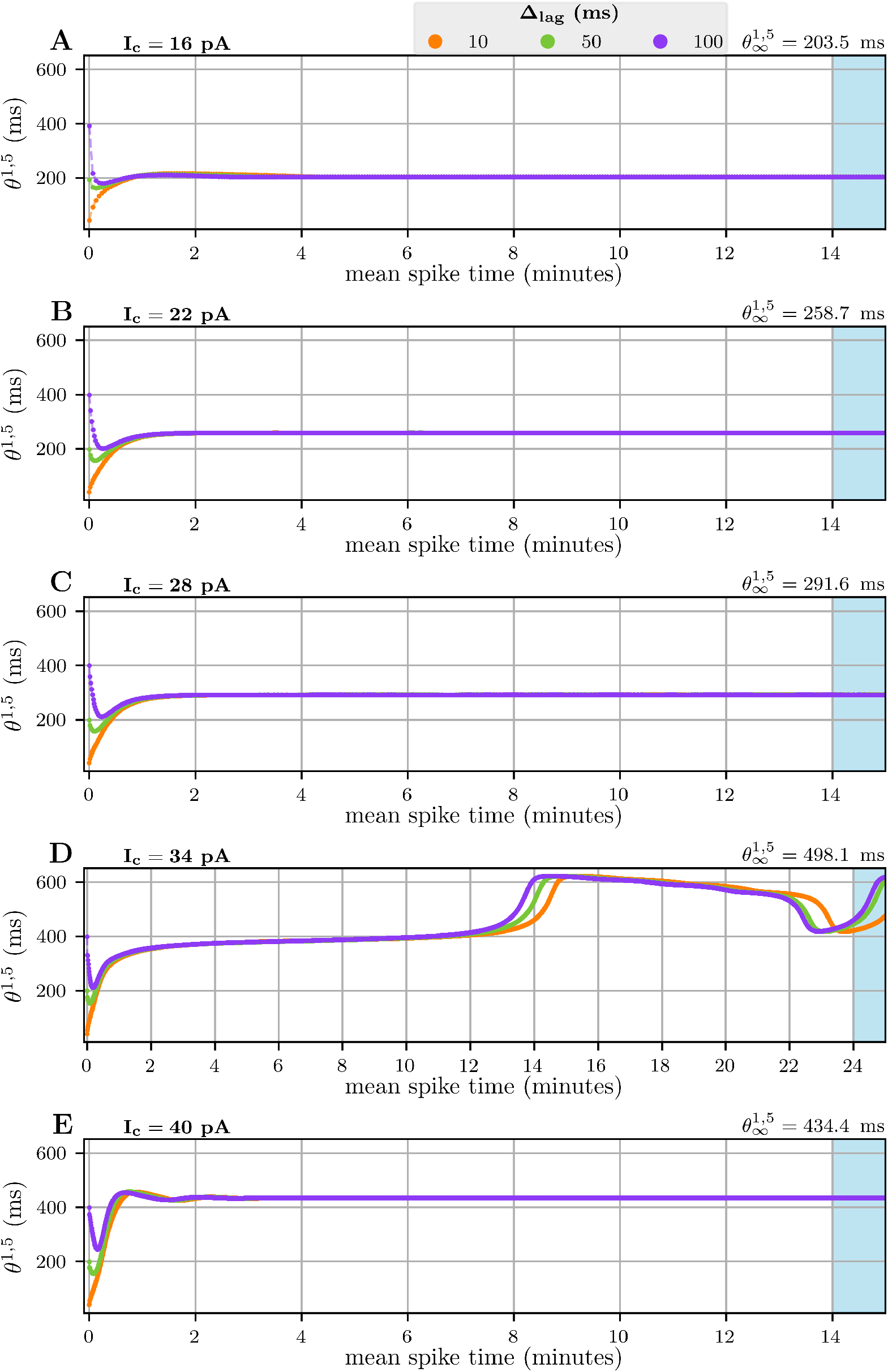
Spike phase difference of five edPR neurons over time. A figure displaying the spike phase *θ*^1,5^ as a function of time. Each panel (**A**–**E**) shows the time evolution of the spike phase difference 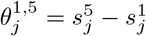, where 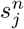 is the *j*’th spike time of the *n*’th neuron. All neurons received a constant stimulus current with stimulus strength *I*_*c*_, starting at time (*n* − 1)Δ_lag_. The stimuli gave population firing rates of 0.35 Hz in **A**, 0.72 Hz in **B**, 1.06 Hz in **C**, 1.43 Hz in **D**, and 1.84 Hz in **E**. Firing rates and converged phases 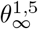 were calculated as the mean value over the last minute (shaded blue area).

**S7 Fig.**
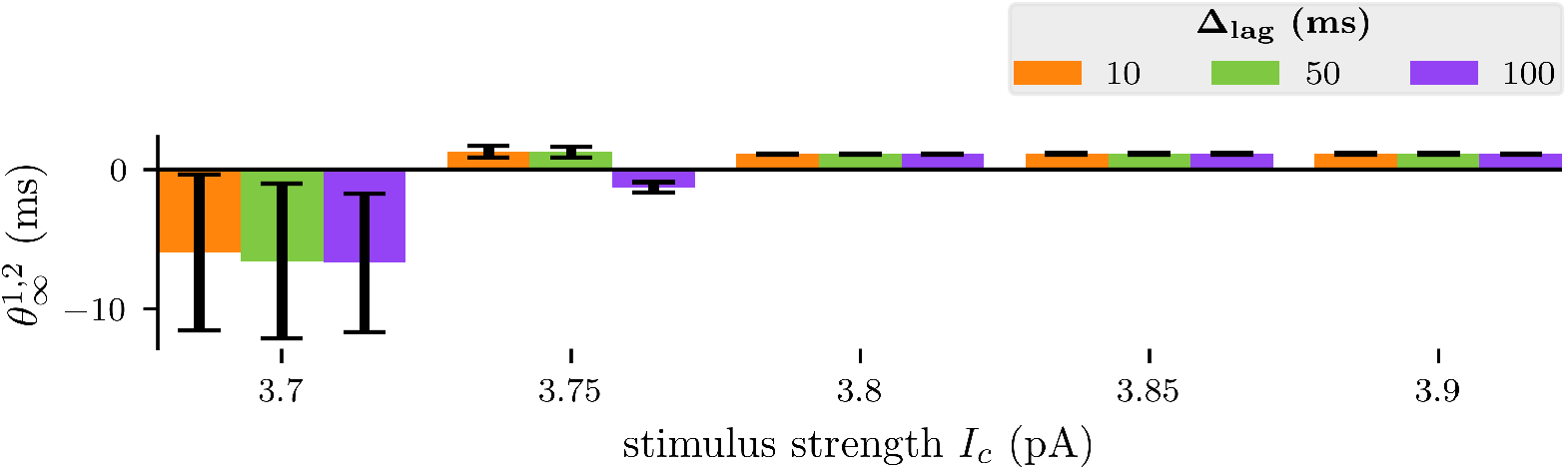
Converged spike phase difference of two edHH neurons. A figure displaying the converged phase differences 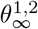 for different stimulus strengths *I*_*c*_ in a system of two edHH neurons. Both neurons received a constant stimulus, each of strength *I*_*c*_ and turned on with a delay of Δ_lag_ from the first to the second neuron. The stimulus strengths *I*_*c*_ were chosen to achieve a similar range of firing rates as the study of 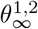 using two edPR neurons in Fig. 8A. The phase differences 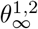 were calculated as the mean value (standard deviation shown with black bar) over the fifth minute to ensure a converged value. Like in the case of edPR (Fig. 8), ephaptic coupling caused spike phase settling among the two edHH neurons. However, the behavior of the two models is qualitatively different. A prominent, albeit not consistent trend for edHH was that the phase difference converged to 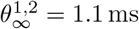, which was consistent only when *I*_*c*_ ≥ 3.8 pA. The trend was independent of choice in both initial delay and stimulus strength. This independence of stimulus strength differs from the results obtained using the edPR model. The edHH model’s spike phase trend shows traditional synchronization, where ephaptic coupling shortens the spike phase. The exception to the spike phase trend of edHH was at the lower firing rates (*I*_*c*_ = 3.7 pA and *I*_*c*_ = 3.75 pA), exhibiting both variation in the spike distance as well as negative mean values. Instead of converging toward a single value, the spike phase 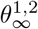 entered a cyclic steady-state (see panels A and B in S8 Fig and a closer look in S9 Fig), causing variation in the mean value 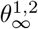. A negative spike phase (*θ*^1,2^ *<* 0) indicates that the two neurons have changed spiking order, meaning the second neuron has *skipped the line*, and now fires *before* the first neuron (illustrated in S10 Fig).

**S8 Fig.**
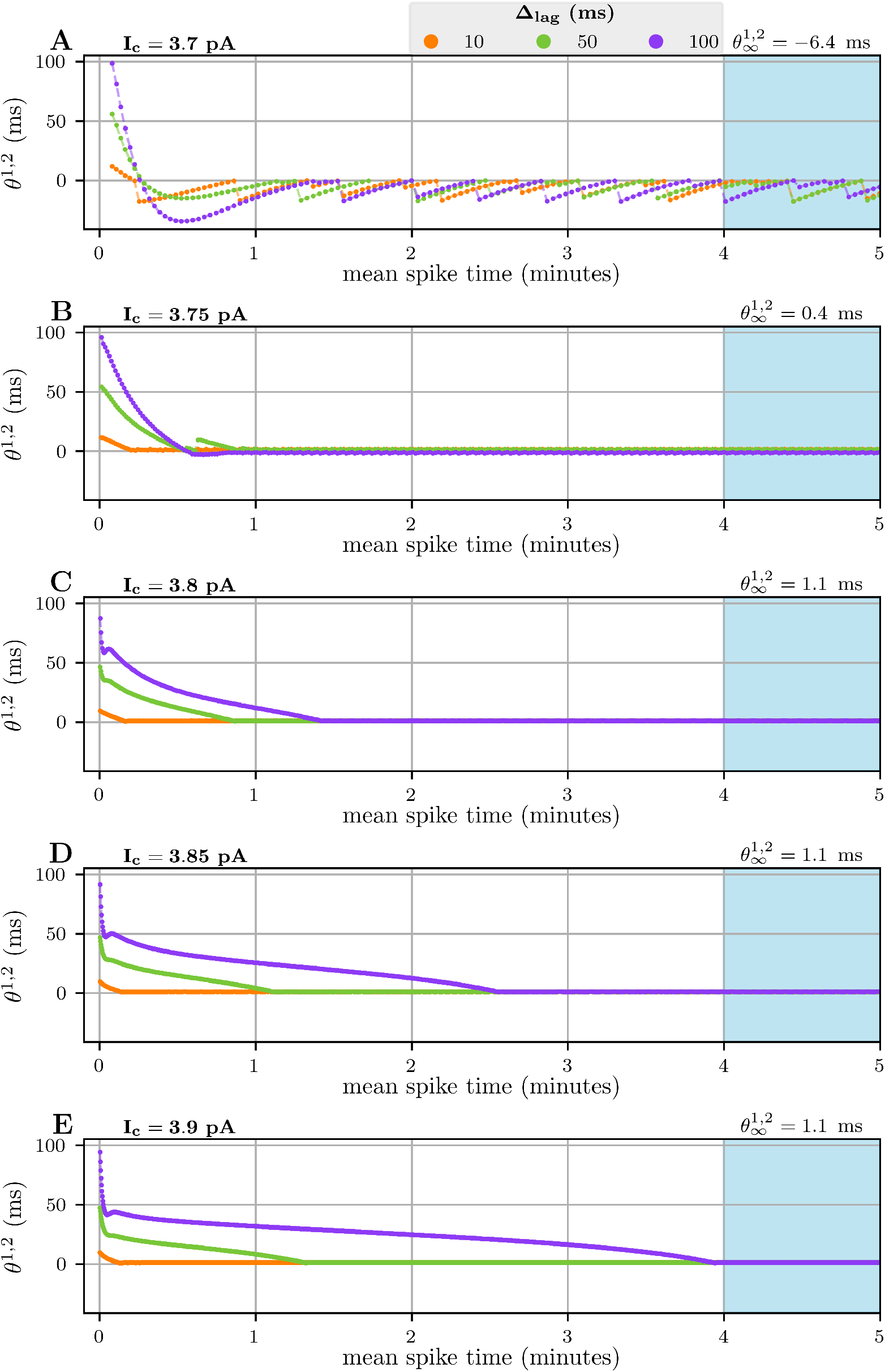
The spike phase difference of two edHH neurons over time. A figure displaying the phase difference *θ*^1,2^ of two edHH neurons as a function of time. Each panel shows the time evolution of 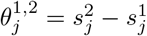, where 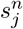 is the *j*’th spike time of the *n*’th neuron. Both neurons received a constant stimulus current with strength *I*_*c*_, with immediate onset for the first neuron and a delayed onset for the second neuron, given by Δ_lag_. The stimuli gave population firing rates of 0.42 Hz in **A**, 0.91 Hz in **B**, 1.35 Hz in **C**, 1.78 Hz in **D**, and 2.17 Hz in **E**. Converged phase differences 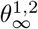 and firing rates were calculated as the mean values over the fifth minute (shaded blue area). While the convergence time was quite consistent for edPR (S5 Fig), it increased with increasing stimulus strength for edHH.

**S9 Fig.**
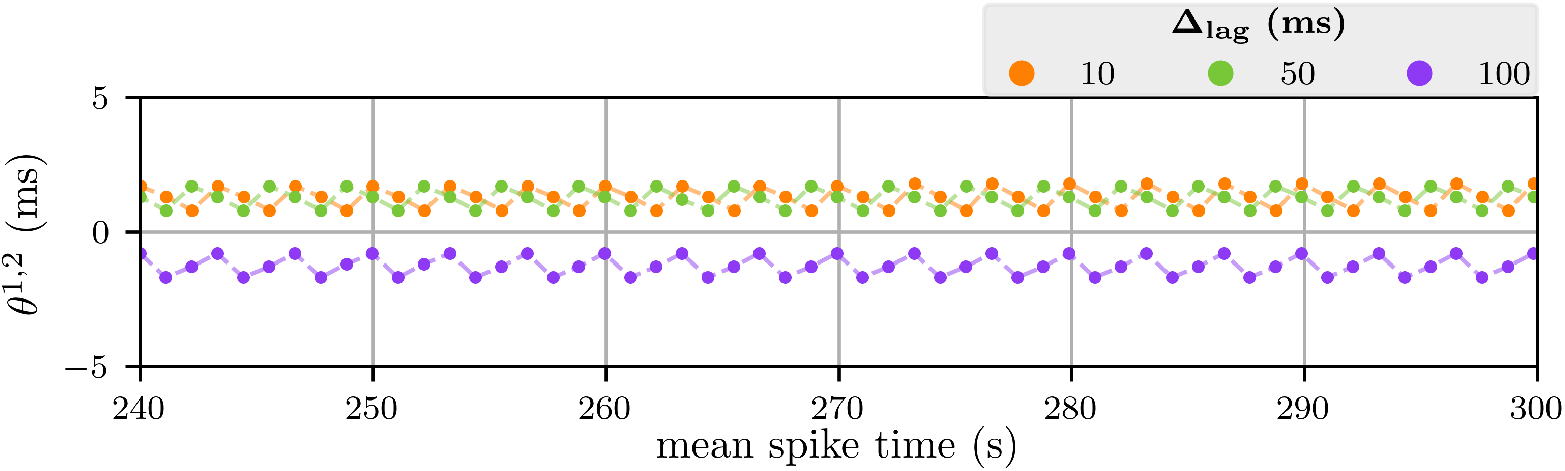
Cyclic spike phase difference of two edHH neurons. The figure provides a close-up of the fifth minute of the phase difference 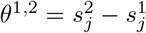 in S8 FigB, where 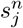 is the *j*’th spike time of the *n*’th neuron. Both neurons received a constant stimulus current with strength *I*_*c*_ = 3.75 pA, with immediate onset for the first neuron and a delayed onset for the second neuron, given by Δ_lag_. In all three cases of Δ_lag_, the phase difference is cyclic with three states.

**S10 Fig.**
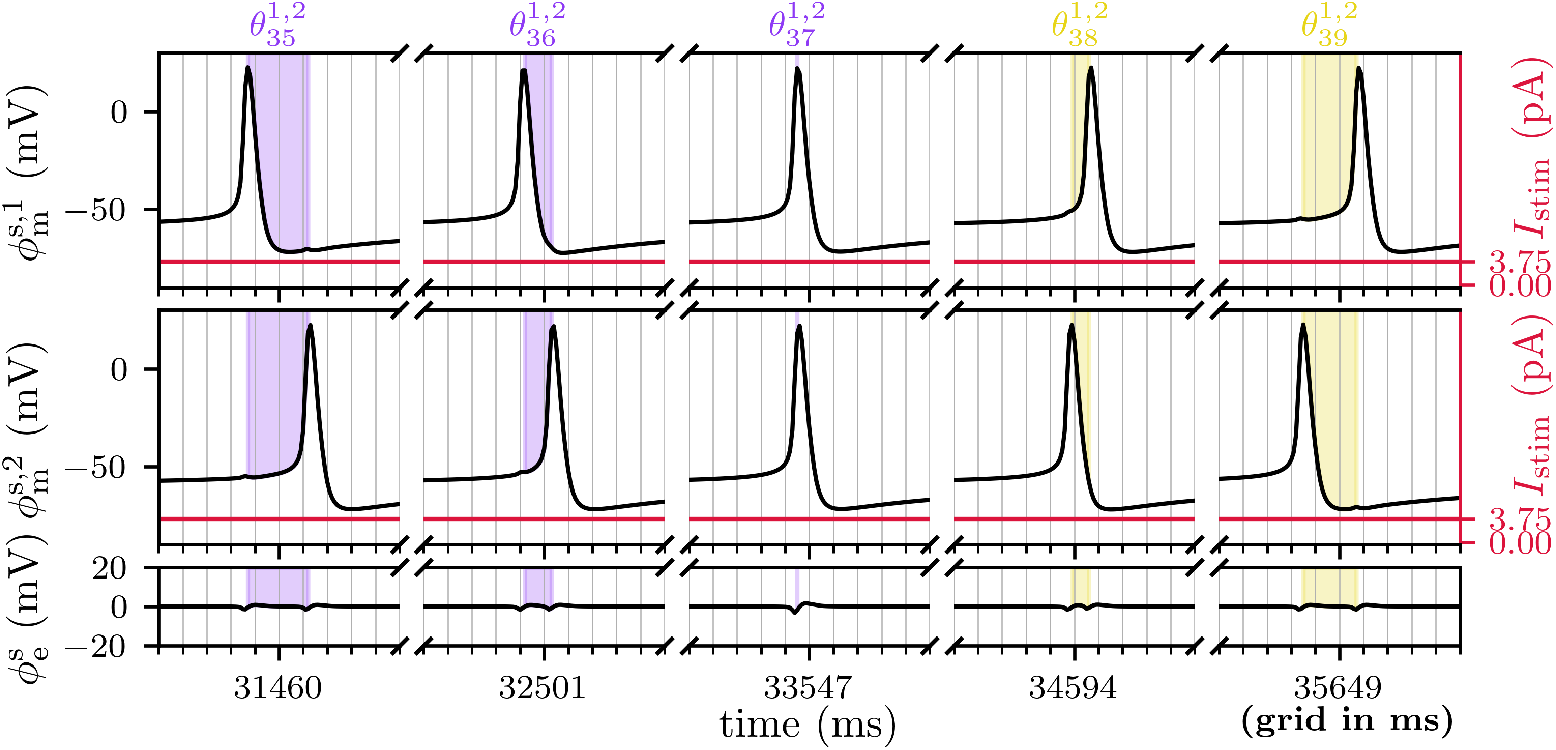
The second neuron *skipping the line*, firing before the first neuron in an edHH simulation. A figure illustrating the change in spiking order between two edHH neurons from the 37th to the 38th spike. Showing the soma membrane potential of the first (upper plot) and second neuron (middle plot), as well as the extracellular potential (lower plot) from the simulation in S8 FigB. Both neurons received a constant stimulus current with immediate onset for the first neuron and a delayed onset for the second neuron, given by Δ_lag_ = 100 ms. All spike phase differences 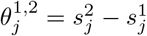, where 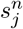 is the *j*’th spike time of the *n*’th neuron, are shaded. Purple shaded areas indicate positive spike phases, while yellow shaded areas indicate negative spike phases. Note that the figures use a broken x-axis, each part spanning 10 ms.

**S11 Fig.**
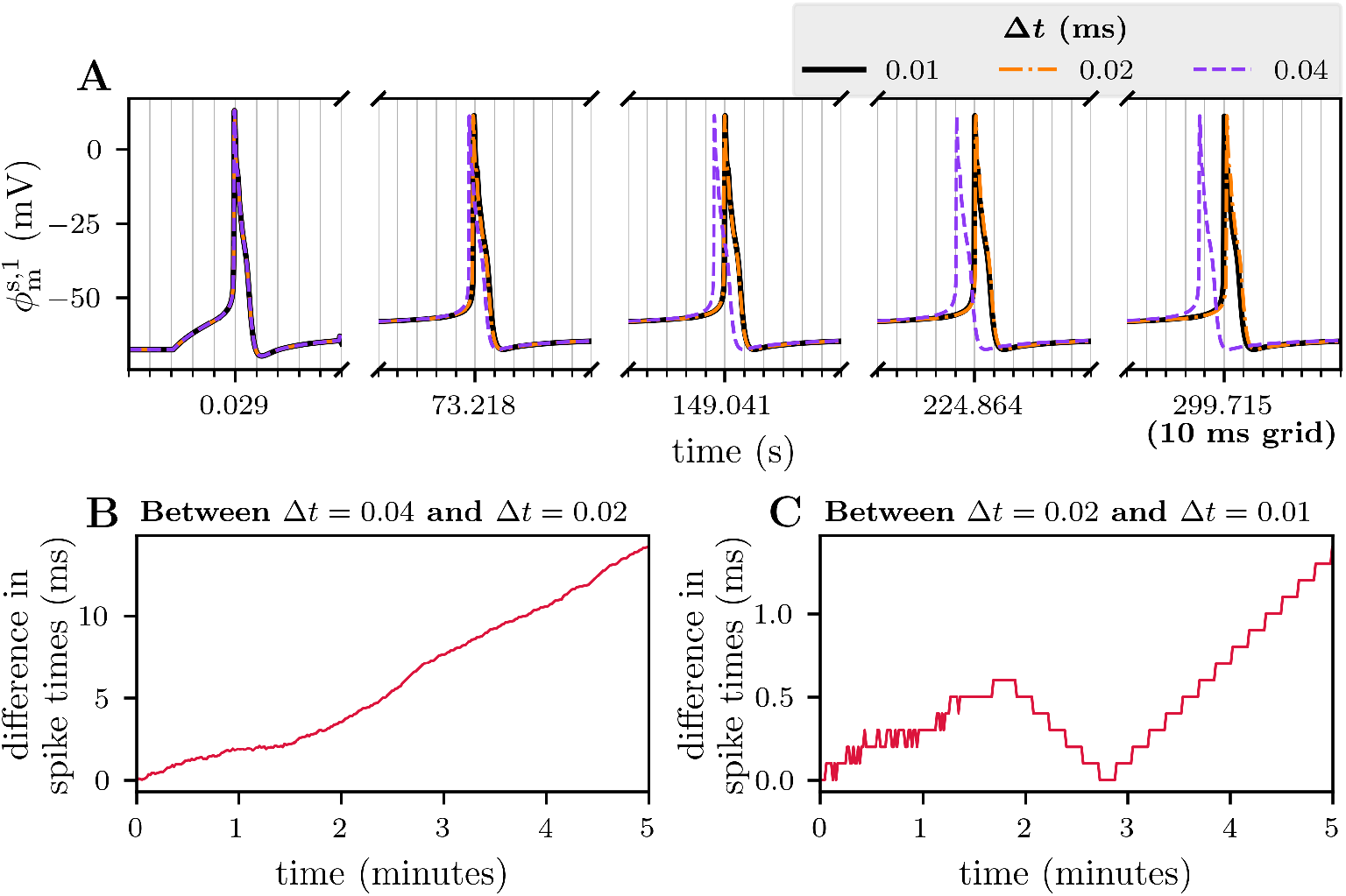
Numerical verification of edPR. A figure comparing spike times using different time step lengths Δ*t* for the numerical integration. The system consisted of two neurons, both receiving a constant stimulus of strength 28 pA. The onset of the first neuron’s stimulus current was immediate, while the second neuron’s stimulus current was turned on after 50 ms. Panel **A** shows the soma membrane potential of the first neuron (note the use of broken axis). Panels **B** and **C** compare the spike times of the first neuron when halving the time step, plotting the absolute difference as a function of time. When reducing the time step from Δ*t* = 0.04 ms to Δ*t* = 0.02 ms (shown in panel **B**), the maximum difference in spike times was 14.2 ms over five minutes of simulated time. Reducing the time step from Δ*t* = 0.02 ms to Δ*t* = 0.01 ms (shown in panel **C**), gave a maximum difference in spike times of 1.4 ms. A time step of Δ*t* = 0.02 ms was considered sufficiently small for the numerical integration of the edPR model.

**S12 Fig.**
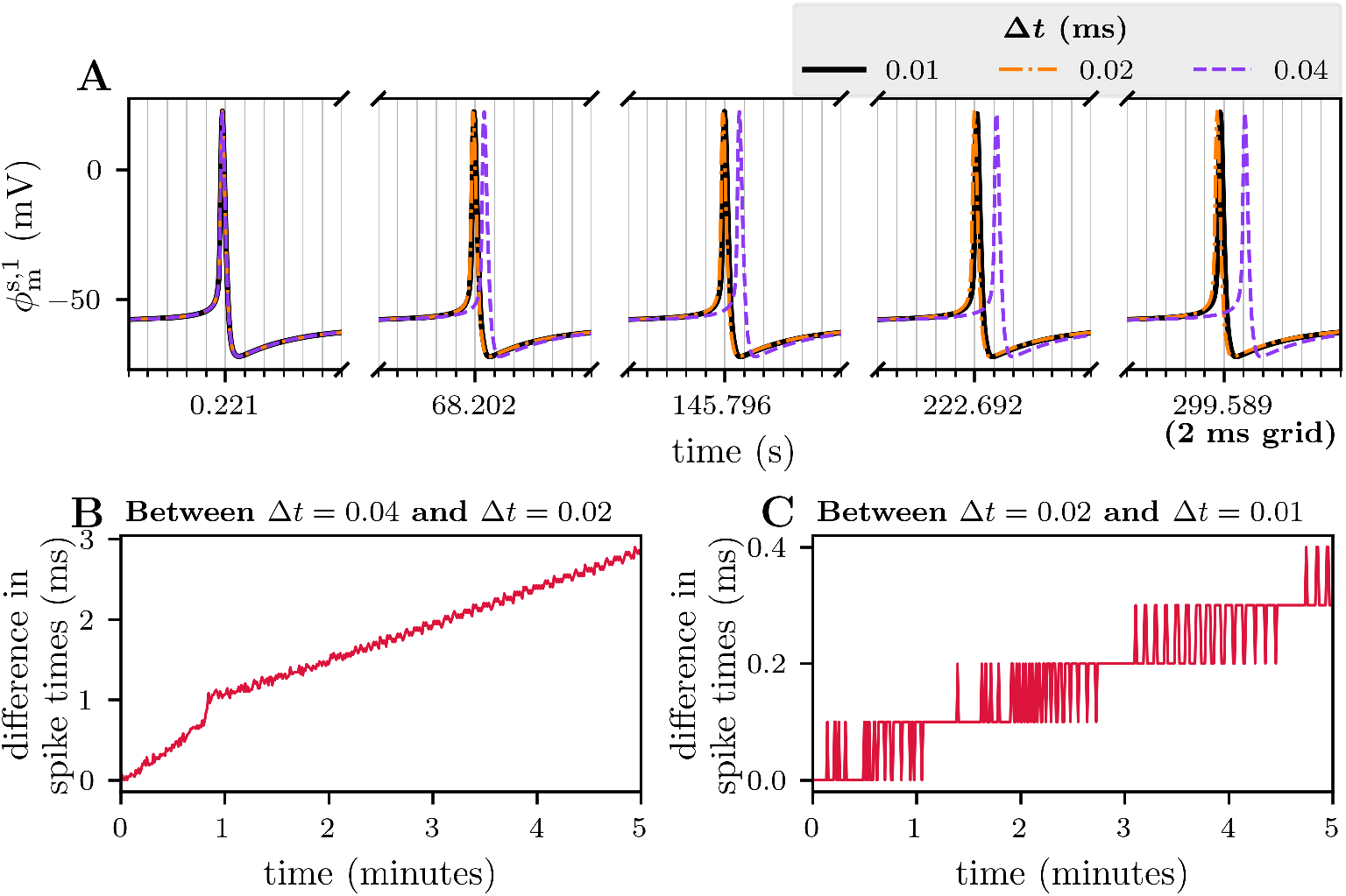
Numerical verification of edHH. A figure comparing spike times using different time step lengths Δ*t* for the numerical integration in an edHH simulation. The system consisted of two neurons, both of which received a constant stimulus of strength 3.8 pA. The onset of the first neuron’s stimulus current was immediate, while the second neuron’s stimulus current was turned on after 50 ms. Panel **A** shows the soma membrane potential of the first neuron (note the use of broken axis). Panels **B** and **C** compare the spike times of the first neuron when halving the time step, plotting the absolute difference as a function of time. When reducing the time step from Δ*t* = 0.04 ms to Δ*t* = 0.02 ms (shown in panel **B**), the maximum difference in spike was 2.9 ms over five minutes of simulated time. Reducing the time step from Δ*t* = 0.02 ms to Δ*t* = 0.01 ms (shown in panel **C**), gave a maximum difference in spike times of 0.4 ms. A time step of Δ*t* = 0.02 ms was considered sufficiently small for the numerical integration of the edHH model.

### S1 Text.

#### Hodgkin-Huxley membrane model

##### S1.1 Dendrite membrane mechanisms

In the electrodiffusive Hodgkin-Huxley (edHH) model, the dendritic membrane is passive, containing only homeostatic mechanisms and leak channels. Their functional forms are the same as for the electrodiffusive Pinsky-Rinzel (edPR) model provided in the main manuscript, and are not repeated in this text. An overview of the edHH dendritic membrane model is provided below.

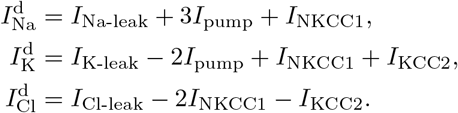

##### S1.2 Soma membrane mechanisms

The soma membrane additionally includes active sodium- and potassium channels, where functionals and maximum conductances are taken from Wei et al. (2014) [15]. The sodium channel is given by

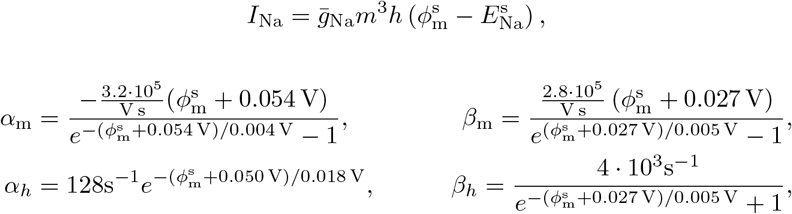

where 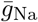 is the maximum conductance, and *m* and *h* are gating variables, evolving in time according to the rate law (described in the main manuscript). The active potassium channel is given by

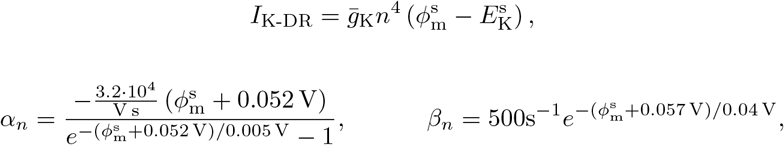

where 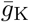 is its maximum conductance, and gating variable *n* also evolves according to the rate law. In sum, the soma membrane channels are given by

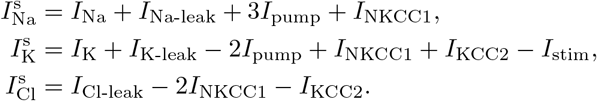

##### S1.3 Model parameters

Only a few of the model parameters differ from the edPR model parameters. These are listed in Table S1.1, and include the membrane capacitance, the sodium leak channel, and the maximum conductances of the active membrane channels. All other parameters are the same as for the edPR model, described in the main manuscript.

Note that the edHH model does not include calcium ions. Therefore, the calcium-dependent membrane mechanisms are not included in the model.

**Table S1.1.**
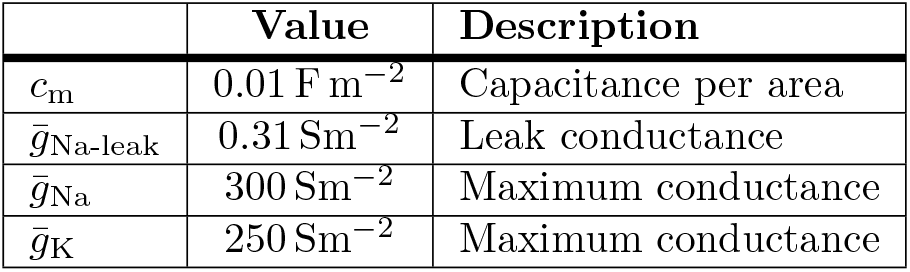
Hodgkin-Huxley membrane parameters. All conductances are taken from [15] except the 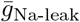, which was tuned to avoid long ramp-up times for the first action potential.

##### S1.4 Calibrating the sodium leak channel

In the initial simulations using the edHH model, we observed long ramp-up times before the first spike, that is, when using a constant weak stimulus current aiming for low firing rates, we saw that the time from the onset of the stimulus until the neurons spiked was exceedingly long compared to the following interspike intervals. Furthermore, we observed a large gap between the resting potential and the membrane potential between action potentials. Therefore, we decided to tune the model to achieve a higher resting potential and thus reduce the inconvenient ramp-up time preceding the first action potential.

In order to raise the resting potential for the edHH model, we decided to strengthen the sodium leak current, although this elevation in potential could be achieved in several ways. Tuning the leak channels provides a simple way to change the resting potential. Of the three leak channels, the sodium channel will have the strongest driving force around the resting potential at physiologically relevant ion concentrations. Tuning the sodium leak conductance should, in absolute values, require the smallest change in a single leak conductance to achieve the desired resting potential.

Varying 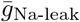, we observed significant changes in the membrane resting potential (several millivolts) while the interspike membrane potential was less affected, staying right above −59 mV and varying by less than a millivolt. After some trial and error, we settled on 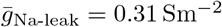, corresponding to an increase of 25.5 %.

##### S1.5 Initial conditions

All steady-state values used as initial conditions are provided in Table S1.2, along with the steady-state values of *ϕ*_m_. The calibration process is described in S3 Text.

**Table S1.2.**
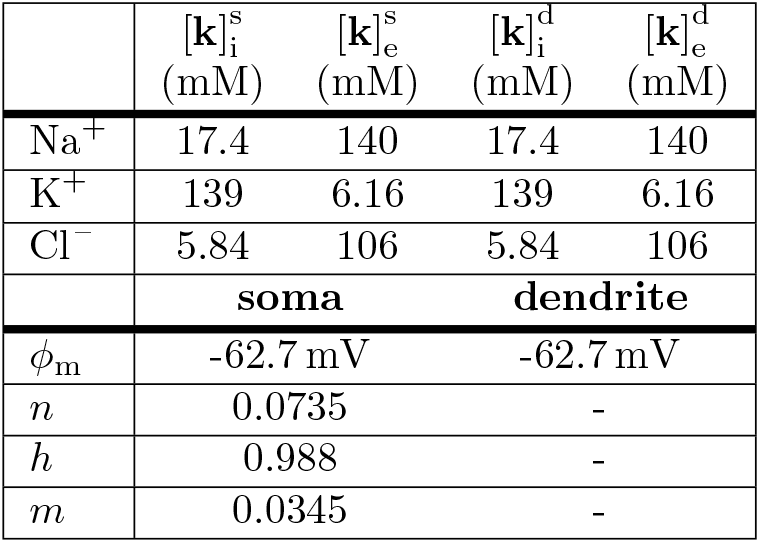
Steady-state values for the edHH model.

#### S2 Text. Optimal ISI length

When the system reaches its ephaptic intrinsic phase preference, the constant distance between the neurons’ spike times indicates that their inter-spike intervals (ISIs) must be identical. Combining this with the observation that the timing of ephaptic interactions affects the ISI length (Section 2.3 in the main text) indicates one of three possibilities: (i) the neurons must fire simultaneously (total synchrony), (ii) the neurons interact ephaptically at the midpoint of their ISIs (which can occur in an idealized two-neuron scenario), or (iii) there must exist multiple times during their ISIs at which ephaptic coupling leads to the same ISI length. We will discuss these possibilities for a system consisting of two neurons, where we assume that the first neuron always spikes before the second, i.e., that 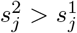, where 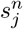 is the *j*’th spike time of the *n*’th neuron.

When the first neuron spikes at spike time 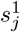, its next spike time 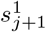, and hence its ISI, is affected by the *other* neurons’ spike time, 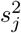. It is not its exact spike time 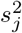 which matters, but rather the distance between spike times, given by

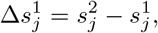

which determines the impact the ephaptic occurrence has on the ISI of the first neuron. In the case of two neurons, the relative spike distance is equivalent to the phase difference, 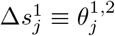. Equivalently, the relative spike distance for the second neuron is given by

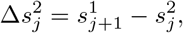

depending on the *next* spike of the first neuron.

The length of the ISI of the *n*’th neuron is a function of this relative spike time distance, as given by

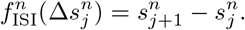

Since the two neurons are identical, they must be equally affected by the ephaptic interactions. This means that the function determining the ISI of the second neuron must be identical to that of the first neuron, i.e., 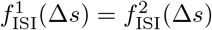, for all spike distances Δ*s*.

When the system has reached its phase preference, the two neurons have the same ISI, meaning that there exist some steady-state spike distances 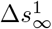 and 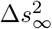 such that

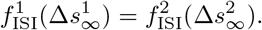

Note that the steady-state spike distance for the first neuron is the same as the phase preference, 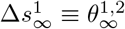. The requirement for the two steady-state spike distances can be achieved in one of two ways.

The steady-state can be achieved if the spike distances are the same, 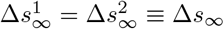. This is achieved either by total synchrony (Δ*s*_∞_ = 0) or the spike times are shifted by exactly half of the ISI, (Δ*s*_∞_ = *f*_ISI_(Δ*s*_∞_)*/*2). However, this is not the case in our studies in Section 2.4.

Since we have that 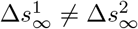, then for 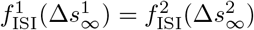 to exist, it requires the function *f*_ISI_ to have at least one extremum. The system containing two, not to mention several spiking neurons, is far more complex compared to the study conducted in Section 2.3. Still, the presence of a minimum in Fig. 5D leads us to believe that a minimum in the ISI length as a function of spike time distances Δ*s* exists also for systems with several spiking neurons. These minima would explain why the system converges into steady-state phase differences, as observed in Section 2.4.

#### S3 Text. Steady-state calibration

A steady-state calibration process was performed for both the electrodiffusive Pinsky-Rinzel (edPR) model and the electrodiffusive Hodgkin-Huxley (edHH) model.

The equilibrium values were obtained by running simulations of a single neuron without injection currents until reaching a steady state. The deviance from a steady state was defined as the norm of the time derivatives, as given by

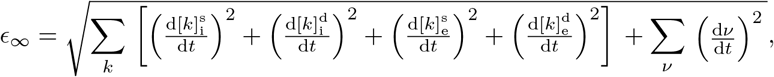

including the derivatives of all ion concentrations [*k*] in all compartments and all gating variables *ν*. We considered the system to be in a steady state when the norm was sufficiently small, i.e.:

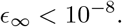

The initial concentrations and gating variables were the same as in the calibration process of Sætra et al. 2020 [17]. Two exceptions apply to the edHH model, where the membrane potential used to find X^−^ was set to *ϕ*_*m*_ = −63 mV and the gating variable *m* was set to its steady-state value, *m* = *m*_∞_(−63 mV). All initial values used in the steady-state calibration process are listed in Table S3.1.

**Table S3.1.** Initial values used to obtain steady-state values. Note that *ϕ*_m_ is not an independent state variable but used initially to determine the concentration of X^−^ .

|  | edHH |  | edPR |  |
| --- | --- | --- | --- | --- |
| | $[k]_i$<br>(mM) | $[k]_e$<br>(mM) | $[k]_i$<br>(mM) | $[k]_e$<br>(mM) |
| $\text{Na}^+$ | 15.0 | 145 | 15.0 | 145 |
| $\text{K}^+$ | 140 | 5.00 | 140 | 5.00 |
| $\text{Cl}^-$ | 4.00 | 110 | 4.00 | 110 |
| $\text{Ca}^{2+}$ | - | - | 0.0100 | 1.10 |
|  | **membrane** |  | **membrane** |  |
| $\phi_m$ | -63.0 mV | | -68.0 mV | |
| $n$ | 0.00100 | | 0.00100 | |
| $h$ | 0.999 | | 0.999 | |
| $m$ | $m_{\infty}(-63 \text{ mV})$ | | - | |
| $s$ | - | | 0.00900 | |
| $c$ | - | | 0.00700 | |
| $q$ | - | | 0.0100 | |
| $z$ | - | | 1.00 | |

Like in the single-neuron edPR model of Sætra et. al [17], the concentration of the generic anion X^−^ was not provided as an initial value but rather calculated to ensure strict electroneutrality. The concentration of X^−^ in each compartment was determined as

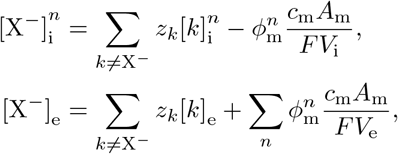

where the concentrations [*k*] for *k* ≠ X^−^ and membrane potentials *ϕ*_m_ are given by the initial conditions.

Both models converged after 6 minutes of simulation time, at which the distances from the steady state were *ϵ*_∞_ = 6 · 10^−9^ for the edPR model and *ϵ*_∞_ = 2 · 10^−9^ for the edHH model. Precise data on the equilibrium values are found in this paper’s GitHub repository.

## References

1. Jefferys J. Nonsynaptic modulation of neuronal activity in the brain: electric currents and extracellular ions. Physiological reviews. 1995;75(4):689–723.

2. Halnes G, Ness TV, Næss S, Hagen E, Pettersen KH, Einevoll GT. Electric brain signals: foundations and applications of biophysical modeling. Cambridge University Press; 2024.

3. Traub RD, Dudek FE, Taylor CP, Knowles WD. Simulation of hippocampal afterdischarges synchronized by electrical interactions. Neuroscience. 1985;14(4):1033–1038. doi:10.1016/0306-4522(85)90274-X.

4. Fröhlich F, McCormick Da. Endogenous electric fields may guide neocortical network activity. Neuron. 2010;67(1):129–43.

5. Clark JW, Plonsey R. A mathematical study of nerve fiber interaction. Biophysical journal. 1970;10(10):937–957.

6. Bokil H, Laaris N, Blinder K, Ennis M, Keller a. Ephaptic interactions in the mammalian olfactory system. The Journal of neuroscience : the official journal of the Society for Neuroscience. 2001;21(20):RC173.

7. Tveito A, Jæger KH, Lines GT, Paszkowski Ł, Sundnes J, Edwards AG, et al. An evaluation of the accuracy of classical models for computing the membrane potential and extracellular potential for neurons. Frontiers in computational neuroscience. 2017;11:27.

8. Shifman AR, Lewis JE. ELFENN: a generalized platform for modeling ephaptic coupling in spiking neuron models. Frontiers in neuroinformatics. 2019;13:35.

9. Goldwyn JH, Rinzel J. Neuronal coupling by endogenous electric fields: Cable theory and applications to coincidence detector neurons in the auditory brain stem. Journal of Neurophysiology. 2016;115(4):2033–2051.

10. Holt G, Koch C. Electrical interactions via the extracellular potential near cell bodies. Journal of computational neuroscience. 1999;6:169–184.

11. Somjen GG. Ions in the brain: normal function, seizures, and stroke. Oxford University Press; 2004.

12. Kager H, Wadman WJ, Somjen GG. Simulated seizures and spreading depression in a neuron model incorporating interstitial space and ion concentrations. Journal of neurophysiology. 2000;84(1):495–512.

13. Ullah G, Cressman JR, Barreto E, Schiff SJ. The influence of sodium and potassium dynamics on excitability, seizures, and the stability of persistent states. II. Network and glial dynamics. Journal of computational neuroscience. 2009;26(2):171–83.

14. Zandt BJ, ten Haken B, van Dijk JG, van Putten MJ. Neural dynamics during anoxia and the “wave of death”. PLOS One. 2011;6(7):e22127.

15. Wei Y, Ullah G, Schiff SJ. Unification of Neuronal Spikes, Seizures, and Spreading Depression. Journal of Neuroscience. 2014;34(35):11733–11743.

16. Hübel N, Schöll E, Dahlem Ma. Bistable dynamics underlying excitability of ion homeostasis in neuron models. PLOS computational biology. 2014;10(5):e1003551.

17. Sætra MJ, Einevoll GT, Halnes G. An electrodiffusive, ion conserving Pinsky-Rinzel model with homeostatic mechanisms. PLOS Computational Biology. 2020;16(4):e1007661. doi:10.1371/journal.pcbi.1007661.

18. Halnes G, Mäki-Marttunen T, Keller D, Pettersen KH, Andreassen OA, Einevoll GT. Effect of ionic diffusion on extracellular potentials in neural tissue. PLOS Computational Biology. 2016;12(11):e1005193.

19. Solbrå A, Bergersen AW, van den Brink J, Malthe-Sørenssen A, Einevoll GT, Halnes G. A Kirchhoff-Nernst-Planck framework for modeling large scale extracellular electrodiffusion surrounding morphologically detailed neurons. PLOS Computational Biology. 2018;14(10):e1006510. doi:10.1371/journal.pcbi.1006510.

20. Sætra MJ, Einevoll GT, Halnes G. An electrodiffusive neuron-extracellular-glia model for exploring the genesis of slow potentials in the brain. PLOS Computational Biology. 2021;17(7):e1008143. doi:10.1371/journal.pcbi.1008143.

21. Anastassiou CA, Koch C. Ephaptic coupling to endogenous electric field activity: Why bother? Current Opinion in Neurobiology. 2015;31:95–103.

22. Pinsky PF, Rinzel J. Intrinsic and network rhythmogenesis in a reduced traub model for CA3 neurons. Journal of Computational Neuroscience. 1994;1(1):39–60.

23. Hodgkin AL, Huxley AF. A quantitative description of membrane current and its application to conduction and excitation in nerve. The Journal of Physiology. 1952;117(4):500–544. doi:10.1113/jphysiol.1952.sp004764.

24. Shafiei M, Jafari S, Parastesh F, Ozer M, Kapitaniak T, Perc M. Time delayed chemical synapses and synchronization in multilayer neuronal networks with ephaptic inter-layer coupling. Communications in Nonlinear Science and Numerical Simulation. 2020;84:105175.

25. Han KS, Guo C, Chen CH, Witter L, Osorno T, Regehr WG. Ephaptic Coupling Promotes Synchronous Firing of Cerebellar Purkinje Cells. Neuron. 2018;100(3):564–578.e3. doi:10.1016/j.neuron.2018.09.018.

26. Rebollo B, Telenczuk B, Navarro-Guzman A, Destexhe A, Sanchez-Vives MV. Modulation of intercolumnar synchronization by endogenous electric fields in cerebral cortex. Science Advances. 2021;7(10):eabc7772. doi:10.1126/sciadv.abc7772.

27. Striebel J, Habibey R, Wendland D, Gehring H, Podoliak E, Pawlick JS, et al. Reproducible Human Neural Circuits Printed with Single-Cell Precision Reveal the Functional Roles of Ephaptic Coupling. ACS Nano. 2025;19(44):38457–38471. doi:10.1021/acsnano.5c11482.

28. Cunha GM, Corso G, Miranda JGV, Dos Santos Lima GZ. Ephaptic entrainment in hybrid neuronal model. Sci Rep. 2022;12(1):1629. doi:10.1038/s41598-022-05343-3.

29. Stacey RG, Hilbert L, Quail T. Computational study of synchrony in fields and microclusters of ephaptically coupled neurons. Journal of Neurophysiology. 2015;113(9):3229–3241. doi:10.1152/jn.00546.2014.

30. Park EH, Barreto E, Gluckman BJ, Schiff SJ, So P. A Model of the Effects of Applied Electric Fields on Neuronal Synchronization. J Comput Neurosci. 2005;19(1):53–70. doi:10.1007/s10827-005-0214-5.

31. Anastassiou CA, Perin R, Markram H, Koch C. Ephaptic coupling of cortical neurons. Nature Neuroscience. 2011;14(2):217–223. doi:10.1038/nn.2727.

32. Øyehaug L, Østby I, Lloyd CM, Omholt SW, Einevoll GT. Dependence of spontaneous neuronal firing and depolarisation block on astroglial membrane transport mechanisms. Journal of Computational Neuroscience. 2012;32(1):147–65.

33. Kofuji P, Newman Ea. Potassium buffering in the central nervous system. Neuroscience. 2004;129(4):1045–56.

34. Larsen BR, Stoica A, MacAulay N. Managing Brain Extracellular K+ during Neuronal Activity: The Physiological Role of the Na+/K+-ATPase Subunit Isoforms. Frontiers in Physiology. 2016;7. doi:10.3389/fphys.2016.00141.

35. Bellot-Saez A, Kékesi O, Morley JW, Buskila Y. Astrocytic modulation of neuronal excitability through K+ spatial buffering. Neuroscience & Biobehavioral Reviews. 2017;77:87–97.

36. Sætra MJ, Mori Y. An electrodiffusive network model with multicompartmental neurons and synaptic connections. PLOS Computational Biology. 2024;20(11):e1012114. doi:10.1371/journal.pcbi.1012114.

37. Ellingsrud AJ, Solbrå A, Einevoll GT, Halnes G, Rognes ME. Finite element simulation of ionic electrodiffusion in cellular geometries. Frontiers in Neuroinformatics. 2020;14:11.

38. Gold C, Henze Da, Koch C, Buzsáki G. On the origin of the extracellular action potential waveform: A modeling study. Journal of neurophysiology. 2006;95(5):3113–28.

39. Syková E, Nicholson C. Diffusion in Brain Extracellular Space. Physiol Rev. 2008;88:1277–1340.

40. Kinney JP, Spacek J, Bartol TM, Bajaj CL, Harris KM, Sejnowski TJ. Extracellular sheets and tunnels modulate glutamate diffusion in hippocampal neuropil. Journal of Comparative Neurology. 2013;521(2):448–464.

41. Halnes G, Østby I, Pettersen KH, Omholt SW, Einevoll GT. Electrodiffusive model for astrocytic and neuronal ion concentration dynamics. PLOS Computational Biology. 2013;9(12):e1003386.

42. Halnes G, Østby I, Pettersen KH, Omholt SW, Einevoll GT. An Electrodiffusive Formalism for Ion Concentration Dynamics in Excitable Cells and the Extracellular Space Surrounding Them. In: Advances in cognitive neurodynamics (IV). Springer Netherlands; 2015. p. 353–360.

43. Halnes G, Mäki-Marttunen T, Pettersen KH, Andreassen OA, Einevoll GT. Ion diffusion may introduce spurious current sources in current-source density (CSD) analysis. Journal of Neurophysiology. 2017;118(1):114–120. doi:10.1152/jn.00976.2016.

44. Ellingsrud AJ, Dukefoss DB, Enger R, Halnes G, Pettersen K, Rognes ME. Validating a computational framework for ionic electrodiffusion with cortical spreading depression as a case study. Eneuro. 2022;9(2).

45. Sætra MJ, Ellingsrud AJ, Rognes ME. Neural activity induces strongly coupled electro-chemo-mechanical interactions and fluid flow in astrocyte networks and extracellular space—A computational study. PLOS Computational Biology. 2023;19(7):e1010996.

46. Herlyng H, Causemann M, Einevoll GT, Ellingsrud AJ, Halnes G, Rognes ME. Modeling and simulation of electrodiffusion in dense reconstructions of cerebral tissue. arXiv preprint arXiv:251203224. 2025;.

47. Qian N, Sejnowski T. An electro-diffusion model for computing membrane potentials and ionic concentrations in branching dendrites, spines and axons. Biological Cybernetics. 1989;15:1–15.

48. Nonner W, Eisenberg B. Electrodiffusion in ionic channels of biological membranes. Journal of Molecular Liquids. 2000;87(2-3):149–162.

49. Mori Y, Peskin C. A numerical method for cellular electrophysiology based on the electrodiffusion equations with internal boundary conditions at membranes. Communications in Applied Mathematics and Computational Science. 2009;4(1):85–134. doi:10.2140/camcos.2009.4.85.

50. Hines ML, Carnevale NT. Neuron: A Tool for Neuroscientists. The Neuroscientist. 2001;7(2):123–135. doi:10.1177/107385840100700207.

51. Hines ML, Davison AP, Muller E. NEURON and Python. Frontiers in Neuroinformatics. 2009;3:1.

52. Park EH, Durand DM. Role of potassium lateral diffusion in non-synaptic epilepsy: a computational study. Journal of theoretical biology. 2006;238(3):666–82.

53. Florence G, Dahlem Ma, Almeida ACG, Bassani JWM, Kurths J. The role of extracellular potassium dynamics in the different stages of ictal bursting and spreading depression: a computational study. Journal of Theoretical Biology. 2009;258(2):219–28.

54. Enger R, Tang W, Vindedal GF, Jensen V, Johannes Helm P, Sprengel R, et al. Dynamics of Ionic Shifts in Cortical Spreading Depression. Cerebral Cortex. 2015;25(11):4469–4476. doi:10.1093/cercor/bhv054.

